# Distinct positions of genetic and oral histories: Perspectives from India

**DOI:** 10.1101/2022.07.06.498959

**Authors:** Arjun Biddanda, Esha Bandyopadhyay, Constanza de la Fuente Castro, David Witonsky, Jose A. Urban Aragon, Nagarjuna Pasupuleti, Hannah M. Moots, Renée Fonseca, Suzanne Freilich, Jovan Stanisavic, Tabitha Willis, Anoushka Menon, Mohammed S. Mustak, Chinnappa Dilip Kodira, Anjaparavanda P. Naren, Mithun Sikdar, Niraj Rai, Maanasa Raghavan

## Abstract

Over the past decade, genomic data has contributed to several insights on global human population histories. These studies have been met both with interest and critically, particularly by populations with oral histories that are records of their past and often reference their origins. While several studies have reported concordance between oral and genetic histories, there is potential for tension that may stem from genetic histories being prioritized or used to confirm community-based knowledge and ethnography, especially if they differ. To investigate the interplay between oral and genetic histories, we focused on the southwestern region of India and analyzed whole-genome sequence data from 158 individuals identifying as Bunt, Kodava, Nair, and Kapla. We supplemented limited anthropological records on these populations with oral history accounts from community members and historical literature, focusing on references to non-local origins such as the ancient Scythians in the case of Bunt, Kodava, and Nair, members of Alexander the Great’s army for the Kodava, and an African-related source for Kapla. We found these populations to be genetically most similar to other Indian populations, with the Kapla more similar to South Indian tribal populations that maximize a genetic ancestry related to Andaman Islanders. We did not find evidence of additional genetic sources in the study populations than those known to have contributed to many other present-day South Asian populations. Our results demonstrate that oral and genetic histories may not always provide consistent accounts of population origins and motivate further community-engaged, multi-disciplinary investigations of non-local origin stories in these communities.

## Introduction

Recent studies have made substantial inroads into characterizing the global genetic diversity of humans. Yet, many populations remain underrepresented in the genetic literature, leaving gaps in our understanding of regional genetic variation, genetic histories, and consequences for human health ^1–3^. While such gaps in the literature contribute to the motivation behind genomic research in underrepresented populations, it is imperative to be mindful of the ethical and socio-political ramifications of the research process for several Indigenous and marginalized peoples ^4–6^. In particular, when investigating genetic histories, conflation between self-identities, derived from a myriad of sources such as oral histories and life experiences, and genetics can lead to harmful consequences for populations ^7–9^. Prioritizing genetic histories over oral histories or using the former to ‘confirm’ the latter can not only be disrespectful to Indigenous knowledge, but also create tension between geneticists, communities, and ethnographers. Instead of attempting to reconcile genetic and oral histories, it is more valuable to recognize the distinct nature and complexities of both forms of knowledge ^10–12^. Unfortunately, oral histories, based on e.g. folk songs and stories, that underpin self-identities and local concepts of ethnogenesis in many populations are dwindling, with limited written records to preserve this unique heritage. All of this calls for efforts to document oral histories and seek out more interdisciplinary ways for geneticists to engage with populations and their cultural histories.

India has a complex history of human migrations and genetic admixture, as well as a rich set of varied socio-cultural practices, all of which have contributed to the extensive cultural and genetic diversity in this region. Previous studies of genetic diversity in India identified population genetic structure that broadly tracks with geography and language ^13–19^. These studies showed that many Indian populations can be modeled along a genetic cline bounded by two statistical constructs: first, Ancestral South Indians (ASI), with genetic components related to Indigenous Andamanese and ancient Iranian farmers and second, Ancestral North Indians (ANI), modeled as a mix of ASI and a genetic component related to Middle-to-Late Bronze Age (MLBA) groups in the central Eurasian steppe region ^15,20,21^. In addition to these larger-scale ancestry gradients, at a finer-scale, genetic structure in India has been impacted by founder events and endogamous practices ^14,21–24^. Despite substantial diversity, characterization of human genetic variation in India has been limited, disproportionate to its proportion of the global population.

This study aims to develop a fine-scale characterization of population structure in Southwest India by recognizing the distinct positions held by inferences from genetic data and oral histories within a community-informed framework. We generated and analyzed whole-genome sequences from individuals identifying as Kodava, Bunt, Nair, and Kapla and, in conjunction with published genome-wide sequences from worldwide populations, investigated genetic histories and population structure in present-day Southwest India (Figure 1). We concurrently present community-engaged documentation of oral histories surveyed by the research team via conversations and observations. The Bunt, Kodava, and Nair have strong self-identities as well as unique cultural traits and oral histories reflected in their origin narratives, some of which are shared between these populations. Compared to these three populations, oral histories of the Kapla, based on anthropological literature and/or community accounts, are less well documented. Available information from communities and historical records reference non-local origins and contacts for all four populations. Here, we considered genetic and oral histories from our investigations to motivate future research on the latter of these populations and, more broadly, on thoughtful ways of bringing together their self identities and genetic histories.

**Figure 1.**
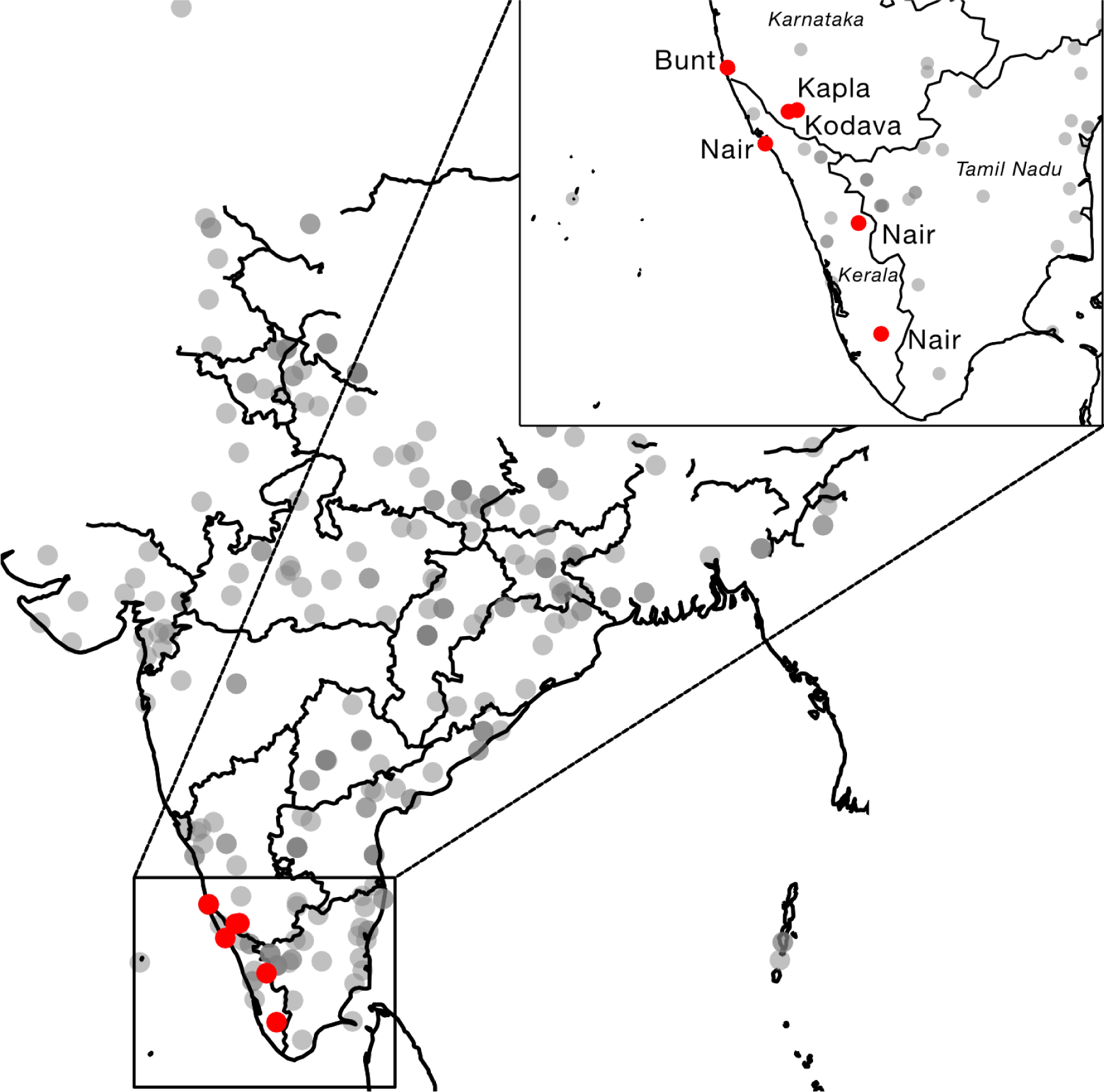
Map of newly sampled populations from this study. Gray points represent regional populations sampled from previous studies ^14,19^.

## Materials and Methods

### Community engagement and sampling

This project was approved by the Mangalore University Institutional Review Board (IRB) (MU-IHEC-2020-4), the Institutional Ethical Committee of Birbal Sahni Institute of Palaeosciences, Lucknow (BSIP/Ethical/2021), and the University of Chicago Biological Sciences Division IRB (IRB18-1572). Community engagement in this study included information sessions in India and the US. Prior to sampling, project aims, methods, and other relevant details were explained to interested community members, and informed consent principles were observed during enrollment in the study. In several locations, local translators, community members, and long-term community contacts joined the research team to facilitate communication with the communities. At these sessions and during subsequent interactions, the research team received valuable input from members and leaders of the study populations on their social organization, cultural traits, oral histories as well as their interest in genetic studies and expectations, which enriched the research process.

Sampling was conducted in India and the US. In India, community contacts and sampling of whole blood were performed in January 2018 in South India in the states of Karnataka, Kerala, and Tamil Nadu. Members of the research team traveled to Kodagu, southern Karnataka, to conduct sampling of Kodava community members (N = 15). The team also conducted sampling of Kapla community members (N = 10), primarily men, who had come down from an isolated mountain settlement close by to work in the Kodagu coffee plantations. The team traveled to Mangalore, southern Karnataka, to conduct sampling of Bunt community members originating in the districts of Udupi and Dakshina Kannada in southern Karnataka and Kasaragod district in northern Kerala (N = 11). Moreover, the team visited four locations in Kerala and Tamil Nadu to conduct sampling of Nair community members (N = 44) with origins in the districts of Pathanamthitta, Palakkad, Kozhikode, and Kannur in Kerala. In the US, the research team was approached by members of the Kodava diaspora to partner on a project investigating their population history. Community engagement was conducted through community representatives who initiated contacts with other community members across the US. Saliva kits, consent forms, and a brief questionnaire were mailed out to interested participants, and 105 participants returned all three to the research team and were enrolled in the study. This dataset was subsequently merged with the data generated from Indian participants, given overlapping aims, with the consent of members of the Kodava_US community.

Results of this study were disseminated to community members of all the study populations, both in India and in the US. Results were returned through a combination of written reports, presentations by the researchers, and a newspaper article, with efforts taken to make the information more accessible to a non-scientific audience. Dissemination was performed both in English and local languages.

### Sample processing and genotype calling

For the Kodava, Nair, Bunt, and Kapla individuals from India, DNA extractions from whole blood were performed in India and were sent to MedGenome Inc, Bangalore, India, for sequencing on Illumina HiSeq X Ten. They were sequenced to an average autosomal depth of 2.5x. For members of the Kodava_US population, saliva samples were collected using the Oragene DNA self-collection kit and extraction was performed using the QIAcube standard protocol in Chicago. The extracts were sequenced to an average autosomal depth of 5.5X on the Illumina NovaSeq 6000 at Novogene, Sacramento, USA in four batches (Table S1). Four samples were re-extracted due to low DNA concentration in the first extraction round and are denoted as ID*_b in Table S1. We re-sequenced three Kodava_US individuals to 77x average depth and two individuals each from Bunt, Kapla, and Nair populations to 30x average depth for variant discovery and for validation of concordance in our genotype calling pipeline. Table S1 summarizes the sequencing results. All reads were subsequently filtered to have a mapping quality score greater than 30 and were aligned to the human reference genome build 37 (hg19) and the revised Cambridge Reference Sequence (rCRS) build 17, for autosomal + Y-chromosome and mtDNA variation, respectively.

Genotypes were called on all high and low coverage samples jointly using the GATK (v4.0) best practices germline short variant discovery workflow on the 8,183,696 sites from the Genome Asia panel ^19^. After this initial phase of genotype calling, the GATK phred-scaled genotype likelihoods were converted to genotype probabilities. IMPUTE2 (v2.3.2) along with the 1000 Genome Phase 3 reference panel were used to perform genotype refinement ^25,26^. After genotype refinement, hard-called genotypes were set to the genotype with the maximum probability and genotypes with a maximum probability less than 0.9 were considered missing. Following genotype calling and refinement, chromosomes were phased using SHAPEIT4 (v4.2.2) along with the 1000 Genome Phase3 reference panel ^27^.

### Merging with external datasets and quality control

We merged our sequence data with publicly available data from the GenomeAsia 100K consortium ^19^ and the Human Genome Diversity Project ^28^. To provide additional context in South Asia, especially to increase representation of Indian populations, we further merged this dataset with genotype data from ^14^. Following these merges, we retained 425,620 SNPs for analysis of population structure.

In order to compare genetic affinities with ancient individuals, we merged samples from the Allen Ancient DNA Resource ^29^, which contains genotypes across ∼1.23 million sites represented on the ‘1240k capture panel’. We also merged the publicly available Human Origins dataset from ^30^, which contains present-day worldwide samples. We used this final merged dataset, consisting of our study samples and five external SNP datasets for all downstream analyses in this manuscript.

Following these merging steps, we filtered samples according to relatedness (up to 2^nd^ degree) using KING and removed samples with > 5% SNP missingness for all downstream population genetic analyses ^31^. We did not apply this filter to data from ancient samples to retain sparse genotyping data for these individuals. Applying these criteria to our data led to the retention of all individuals as described in the *Sampling and community engagement* section, except for the Kodava_US, from which we excluded 27 genomes for a final sample number of 78 from the 105 individuals sampled initially. The applied filters also retained 425,620 unique autosomal SNPs on which all subsequent analyses were performed.

### Principal Component Analysis and ADMIXTURE

We performed a principal component analysis (PCA) using smartpca v7.3.0 ^32,33^. We filtered the merged datasets according to linkage disequilibrium using PLINK 1.9 [`--indep-pairwise 200 25 0.4`] ^34^. We chose to plot only population median locations in PCA-space (as gray text labels) of populations from external datasets for ease of visualization, with populations exclusive to this study represented by larger dots of the same color.

This same set of LD-pruned variants was used to run ADMIXTURE v1.30, with *K* = 6 to 11^35^. We restricted visualization of clustering results in the main text figure to representative populations from each geographic region with at least 8 samples.

### Admixture history using Treemix

We modeled the relationship between the study populations in Southwest India and a subset of present-day and ancient Eurasian groups (Table S2) using Treemix ^36^. Positions were filtered by missingness and LD-pruned with PLINK 1.9 as for PCA. The analysis was run, estimating up to 10 migration edges with Mbuti as the outgroup.

### Ancestry estimation using *f*-statistics

For calculating *f*-statistics defined in ^37^, we used the ADMIXTOOLS v7.0.2 software. The first statistic we estimated was the outgroup *f_3_*-statistic, which measures the shared drift or genetic similarity between populations (A, B) relative to an outgroup population (O) and was calculated using the `qp3pop` program. We used the Mbuti population as the outgroup, unless otherwise noted ^28^.

To model the study populations (*Study*) as mixtures of ANI and ASI ancestries, we used the *f_4_-*ratio test ^20^. We measured the proportion of Central Steppe MLBA (α) genetic ancestry via the ratio (see Figure S1) ^15^:

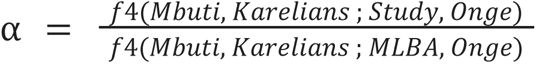

We selected populations to include in the above topologies based on the scheme in ^21^. Briefly, we ran a *D*-statistic using the `qpDstat` program of the form *D(Central_Steppe_MLBA, Iran_GanjDareh; H3; Ethiopian4500)* to evaluate the degree of allele sharing between Central Steppe MLBA and H3. As H3, we selected a large array of present-day populations from Eurasia. We identified the population with the highest value in this test (Karelia) as the closest population to Central Steppe MLBA (Figure S2).

We computed allele sharing of sub-clusters of Kapla based on PCA, Kapla_A and Kapla_B, with different present-day populations (H3) from Europe, the Middle East, and South Asia on the ANI-ASI cline through the topology *D(Kapla_B, Kapla_A; H3, Mbuti)* ^37^, using the `qpDstat` program. For all computed *D*-statistics, we used the (`f4mode: NO, printsd: YES`) options in `qpDstat.`

### Estimation of ancestry proportions based on qpAdm

To model the ancestry proportions of the study and other select Indian populations for comparison, we used qpAdm ^38,39^ implemented in ADMIXTOOLS v7.0.2 software with (`allsnps: YES, oldallsnpsmode: NO, inbreed: NO, details: YES`). As many present-day South Asians on the ANI-ASI cline can be modeled as a mix of genetic ancestries related to the Indus Periphery Cline, Onge, and/or Central Steppe populations from the Middle and Late Bronze Age ^15^, we considered these as our genetic sources for the target populations included in the analysis (Table S3). For the outgroups, we used the list outlined in the proximal model of present-day South Asians by ^15^ (Table S3). Leveraging oral history knowledge, we also ran tests including as a fourth source published genome-wide data from various Scythian groups from ^40–42^ for the two Kodava groups, Nair, and Bunt as targets. As proxy for genetic diversity contemporaneous to Alexander the Great’s reign and may have contributed to his army, we included as a fourth source Classical and Hellenistic period groups from Greece and Macedonia, a Late Bronze Age group from Armenia, and Iron Age groups from Turkmenistan, Iran, Pakistan, and Anatolia ^43,44^ for the two Kodava groups as targets (Table S3). For these tests, we modified the outgroup list from ^15^ used in the three source model by additionally including proximal outgroups to the Scythian (Israel_Natufian, Altaian) and Greek and Macedonian Classical and Hellenistic period (CHG, Italy_Sicily_LBA, Jordan_PPN) source groups.

### Admixture dates based on LD decay in ALDER

To estimate the timing of ANI-related genetic admixture in the study populations, we used the weighted admixture linkage disequilibrium decay method in ALDER ^45^. The true ancestral populations for South Asian populations are unknown, so we followed the approach originally proposed in Moorjani et al. ^15,21^. Briefly, this modified approach approximates the spectrum of ANI-ASI admixture using PCA-derived SNP-weights to capture ancestry-associated differences in allele frequency without requiring explicitly labeled source populations. Following previous studies, we included primarily Dravidian- and Indo-European-speaking populations in India that fall on the ANI-ASI cline on the PCA and have greater than five individuals, and Basque individuals from Europe as an anchor for Western Eurasian ancestry when constructing the SNP-weights (Table S4) ^21^. We estimated the timing of admixture in generations and years, assuming a generation time of 28 years ^46^. Individuals from the focal test population for admixture were excluded when computing PC-based weights to limit biases. For this reason, we also excluded the Coorghi from the reference panel and the test group due to historical connections between the names ‘Coorg’ and ‘Kodagu’, referring to the same region in Southwest India. Based on historical accounts, Coorghi and Kodava may be the same or closely-related communities. Unless otherwise stated, we use Z-score >= 2 (p-value < 0.05) across ALDER tests as indicative of a significant signal of admixture ^21,45^.

### Haplotype-based Estimation of population structure using fineStructure

We implemented a haplotype-based approach using the software ChromoPainter and fineSTRUCTURE ^47^. Briefly, this approach estimates a co-ancestry matrix based on haplotype sharing under a haplotype-copying model (ChromoPainter), which can be used to identify population structure through clustering (fineSTRUCTURE). We first estimated the effective population size (Ne) and the mutation parameter θ in a subset of chromosomes (1, 5, 10, and 20) with 10 expectation-maximization iterations. The average value for both parameters was estimated across chromosomes and used in subsequent runs (Ne = 401.352, θ = 0.0002). We standardized to five individuals per population, filtering out groups with a sample size below this threshold and randomly sampling five individuals from groups with larger sample size. The co-ancestry matrix obtained from ChromoPainter was used for further inferences in fineSTRUCTURE. The analysis was run at different levels of population composition; the first included all Eurasian populations, followed by South Asia-specific runs. The per-locus copying probabilities across all haplotypes in a population under the maximum-likelihood parameters were used to estimate the median per-locus copying probabilities.

### mtDNA and Y-chromosome analyses

The alignment files for each sample were used to retrieve reads mapping specifically to the mitochondrial genome (mtDNA) using `samtools` ^48^. Variants from mtDNA reads for each sample were called against the revised Cambridge Reference Sequence (rCRS) using `bcftools mpileup` and `bcftools call`, requiring a mapping quality of 30 and base quality of 20. The 158 resulting mtDNA variant calls were used as input for assigning mtDNA haplogroups using the program Haplogrep2 with PhyloTree mtDNA tree Build 17 ^49,50^. The final haplogroup assignments reported in the study were based on the highest quality score and rank assigned by Haplogrep2.

For assigning Y-chromosome haplogroups, we first extracted reads mapping to the Y-chromosome using `samtools`. Variants from Y-chromosome reads were then called using `bcftools mpileup` and `bcftools call`, requiring a mapping quality of 30 and base quality of 20 (`-d 2000 -m 3 -EQ 20 -q 30`). The 100 resulting variant calls were then used to call haplogroups using Yhaplo ^51^ (Table S12).

To investigate the influence of matrilocality and patrilocality onuni-parentally inherited markers, we calculated haplotype and nucleotide diversity on the mitochondrial DNA ^52,53^. We estimated standard deviations of haplotype diversity within populations using 100 bootstrap samples and used one-sided t-tests to evaluate differences between haplotype diversity.

### Pseudo-haploidization of whole-genome sequencing data

We replicated select analyses with pseudo-haploid calls to provide confirmation of many of our population genetic results in light of the lower depth of coverage for many of the sequenced individuals. The alignment files were used to generate pseudo-haploid calls for the Kodava, Kodava_US, Nair, Bunt, and Kapla individuals. We first used `samtools mpileup` with the flags (`-Q 30 -q 20 -R -B`) to generate a mpileup output format for the individuals’ alignment files. The resulting mpileup file was then used as input for `pileupCaller` from the SequenceTools package (https://github.com/stschiff/sequenceTools).

### Estimation of IBD Scores and Founder Events

To measure the extent of endogamy in each population, we evaluated the IBD score, which is a measure of recent strength of consanguinity ^14^. The IBD score is defined as the sum of IBD between 3 cM and 20 cM detected between individuals of the same population, divided by 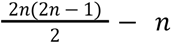, where n is the number of individuals. We used GERMLINE2 to call IBD segments with the flag `-m 3` to only consider IBD segments at least 3 cM in length, using the deCODE genetic map coordinates for genome build hg19 ^54^. We also removed individuals related up to second degree using KING ^31^ and removed IBD segments > 20 cM ^14^. In the original application of the IBD score ^14^, the authors normalized to individuals of Finnish and Ashkenazi Jewish ancestry as canonical examples of founder populations with a higher recessive disease burden. However, we left the IBD scores as raw values, so we only interpreted the results relatively between Indian populations we tested against.

To characterize founder events, we used ASCEND, with default standard parameters for present-day populations ^55^. We selected Mbuti as an outgroup, under the assumption that this population has limited shared recent demography with the target populations. We reported a founder event to be significant if four criteria were met ^55^: (i) the 95% CIs of the estimated founder age and intensity did not include 0, (ii) the estimated founder age was < 200 generations before present and its associated SE was < 50 generations, (iii) the estimated founder intensity was > 0.5%, and (iv) the normalized root-mean-square deviation was < 0.29.

### Runs of Homozygosity

We estimated runs of homozygosity (ROH) using PLINK 1.9, using the parameters: `--homozyg-window-snp 50 --homozyg-snp 50 --homozyg-kb 1500 --homozyg-gap 1000 --homozyg-density 50 --homozyg-window-missing 5 --homozyg-window-het 1` ^56,57^. To increase SNP density, we limited our analysis to a panel including only whole-genome sequencing data from GenomeAsia 100K ^19^ and the Human Genome Diversity Project ^28^. After filtering positions with more than 0.5% missing data and MAF of 5%, the final database retained 3,288,336 variants.

### Novel Variant Discovery

To perform variant discovery, we relied on the nine high-coverage whole genome sequencing (WGS) samples (coverage > 30x) from four populations. In total, we had three Kodava_US, two Bunt, two Kapla, and two Nair individuals in this smaller set of individuals with WGS data. Alignment to GrCh37 (hg19) was performed using bwa-mem ^58^ (Table S1). We called genotypes within these individuals using GATK v.3.0 best practices and filtered to SNPs and indels with a phred-scaled quality score of > 30 to carry forward for variant annotation.

For variant annotation, we used Ensembl Variant Effect Predictor (VEP) v105.0 with custom annotations derived from GnomAD genomes site-level VCF files and GenomeAsia site-level VCF files. In addition, we used the `--af_gnomad --af_1kg` flags to detect whether the alleles were found in the 1000 Genomes project or in GnomAD exomes. Therefore, our definition of novel variants only includes variants that were not found in any of the aforementioned datasets. When assessing variant effects, we used the broad categories provided by VEP (MODIFIER, LOW, MODERATE, HIGH).

## Results

### Survey of population oral histories relating to origin stories

While the Kodava, Bunt, and Nair have strong self-identities and associated oral histories, relatively little is documented in the ethnographic literature ^59–63^. These populations are all speakers of Dravidian languages. Moreover, there are notable overlaps in these populations’ traditions and cultural traits, which may stem from geographical proximity and historical contacts that are also referenced in these populations’ oral histories ^63^. These include historical ‘warrior’ status designations/identities and matrilineal descent in the Bunt and Nair, with the latter additionally practicing matrilocality in the recent past. Moreover, the Nair have a complex and longstanding social system that consists of several subgroups ^64^. Some of these socio-cultural characteristics are noted in the anthropological literature and were additionally shared by participants and community representatives with the research team during fieldwork ^65,66^. Similarly, there are anecdotal accounts, for instance, noted in blogs maintained by community members and relayed to members of the research team during sampling and subsequent population interactions, of phenotypic characteristics that members of these populations draw on to support possible non-local origins. From our interactions with these populations, these narratives are used by population members to contemplate unique phenotypes and customs such as music, diet, religious affiliation, and traditional wear that set them apart from neighboring populations. Since the exact mode and timing of past population contacts may not always be explicitly stated in oral histories, it is unknown whether the nature of the non-local population interactions that they invoke was socio-economic, genetic, or both.

One link to non-local populations shared across the oral histories of the Bunt, Kodava, and Nair is to the Scythians, ancient nomadic people inhabiting the Eurasian Steppe during the Iron Age. In addition, the Kodava have oral histories suggesting links to the Kurdish Barzani tribe through the latter’s involvement in Alexander the Great’s campaign into India. The following is noted in ^67^: “*In those days, when the army advanced, their families of fighting men too moved behind them, as camp followers. After Alexander turned back, some tribes in his army who had no energy to get back to their homeland, stayed back in India.…..Our ancestors* [Kodava, are] *believed to have taken a southernly route along with Western Ghats in search of better prospects* [and] *eventually settled in Kodagu (Coorg) which was then an unnamed, inhospitable and extremely rugged hilly region*”. Height and nose shape were cited during our interactions with Kodava_US community members as examples of phenotypic characteristics that contribute to the physical distinction between the Kodava and neighboring populations and as evidence of non-local origin.

Field observations and interviews by members of the research team suggested that the Kapla are socio-culturally different from the other populations sampled in this study. To our knowledge, there is very little documentation on the Kapla in the anthropological literature. The Kapla population lives geographically close to the Kodava in the Kodagu region of southern Karnataka. They still practice a form of hunting and gathering, but their subsistence is transitioning slowly with increased contacts with the Kodava, on whose coffee plantations Kapla men work during the harvesting season. They speak a mixture of Tulu and Kodava languages, both members of the Dravidian language family. Historical records suggest ^68^: “*The Kaplas, who live near Nalkanad palace seem to be mixed descendants of the Siddis - the Coorg Rajahs’ Ethiopian bodyguard - as their features resemble the Ethiopian type. They have landed property of their own near the palace, given by the Rajahs, and work also as day laborers with the Coorgs. Their number consists of only 15 families*”. Siddis are historical migrants from Africa who live in India and Pakistan ^69^. Anthropological information gathered by the research team from long-term community contacts suggests that once the Kapla settled in their present location in Kodagu, they were isolated from neighboring populations. Furthermore, the Kapla follow a patriarchal and patrilocal system, with a preference towards marriages involving MBD (mother’s brother’s daughter) and junior sororate marriage practices.

### Broad-scale population structure in Southwest India

We used principal component analysis (PCA) to explore the broader population structure within South Asia, with a particular focus on Southwest India. The main axes of genetic variation highlight the ANI-ASI genetic cline (Figure 2A, B, Figure S3). The Kodava, Kodava_US, Bunt, and Nair populations are placed in PCA-space close to other populations that are geographically proximal (e.g., Iyer, Iyangar, and individuals from Urban Bangalore from ^14,19^). This pattern was also captured by genetic ancestry clusters from ADMIXTURE where, at K = 7, the Kodava, Kodava_US, Bunt, and Nair shared genetic components (colored pink and gray in Figure 2C, Figure S4, S5) with many other populations from South Asia. Additionally, they displayed components maximized in the Kalash (colored orange) and in Europe (colored red) that were also observed in populations from Central Asia and the Middle East as well as in most North Indian and some South Indian populations. Notably, the proportion of all these components in the Bunt, Kodava, Kodava_US, and Nair were in line with the proportions displayed by geographical neighbors such as Iyer and individuals from Urban Bangalore and Urban Chennai from ^19^. Furthermore, we did not detect any structure within the Nair despite sampling broadly across the state of Kerala or within the Kodava_US donors who, while being recent migrants to the US, originate from various locations within the Kodagu district in southern Karnataka (Figure 1). Based on these analyses, we also did not observe any notable differences in genetic ancestry between Kodava sampled in India and the Kodava sampled in the US (Kodava_US). These results are consistent with the diaspora being too recent for appreciable genetic differentiation and that recent migrants to the US are a representative random sample of the Indian population with respect to genetic ancestry.

**Figure 2.**
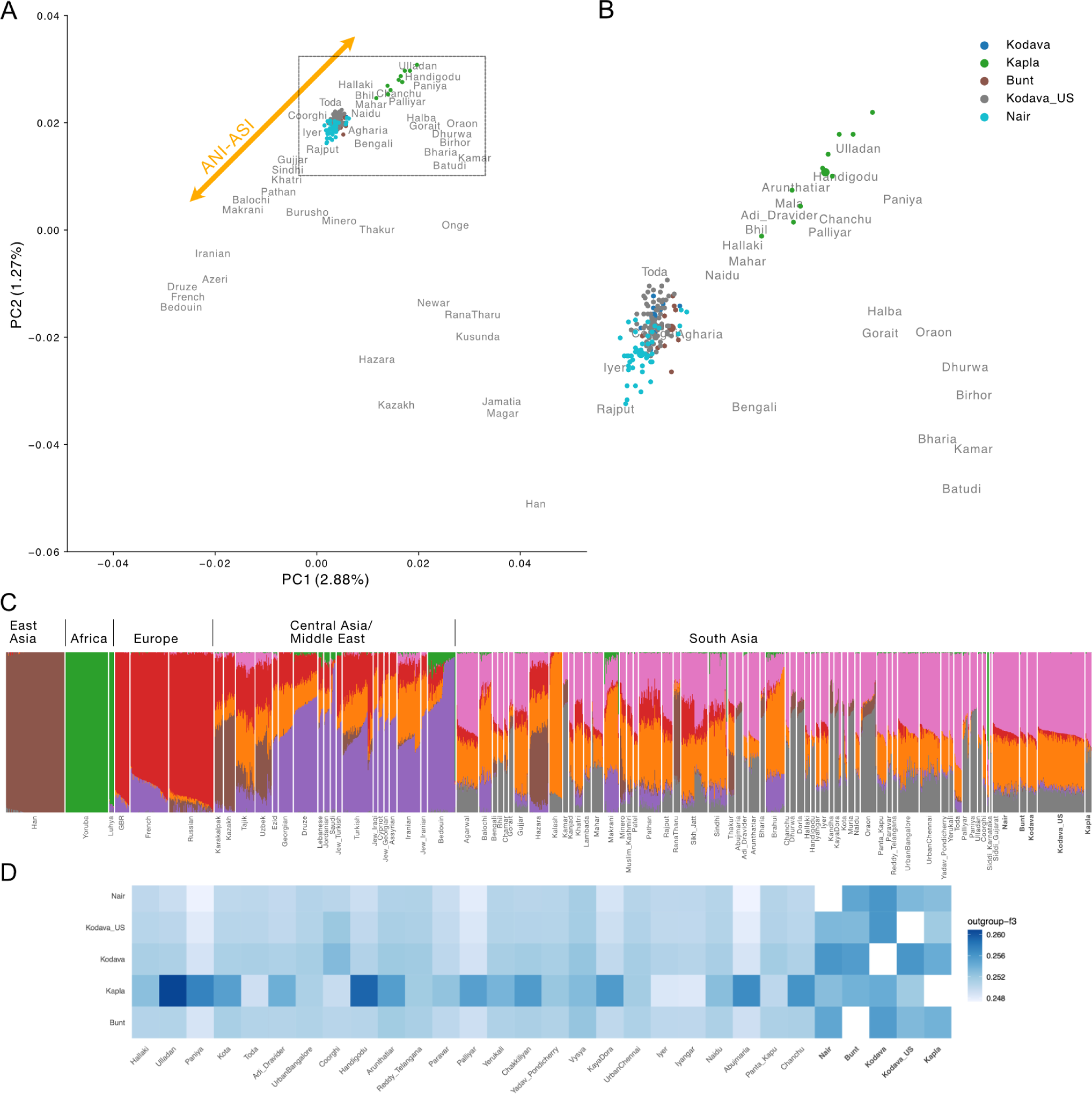
(A) Principal component analysis of merged dataset with Human Origins samples (see Figure S3 for replication with pseudo-haploid calls), with (B) representing a zoomed inset focusing on the study populations and their position on the ANI-ASI genetic ancestry cline. Population labels in gray represent the mean location of all samples in PCA. For populations sampled specifically in this study, mean locations are represented as larger circles of the same color. (C) ADMIXTURE results (*K=7)* showcasing genetic similarity across Southwest Indian populations in relation to African, East Asian, Central Asian, Middle Eastern, and other South Asian populations. We removed some population labels for visual clarity (see Figure S4 for a full set of labels and Figure S5 for replication with pseudo-haploid calls). Populations sampled specifically in this study are represented using bolded population labels. (D) Outgroup *f_3_*-statistics results for the study populations relative to other South Indian populations. See Figure S7 for an extended panel consisting of worldwide populations and Figure S8 for replication with pseudo-haploid calls.

On the other hand, the Kapla showed genetic similarity to populations with higher ASI-related genetic ancestry from South India, such as the Ulladan and Paniya, displaying pink and gray components found in most South Asians but substantially lower orange component maximized in the Kalash and found in populations with higher ANI-related genetic ancestry (Figure 2A, B, C). In addition to being genetically differentiated from the other populations sampled in this study, the Kapla also exhibited greater spread on the PCA than other South Indian populations maximizing ASI ancestry, such as Ulladan and Paniya (Figure 2B). Finally, we found no qualitative support for substantial genetic similarity between the Kapla and the Siddi or tested African individuals (Figure 2B, Figure S6).

We evaluated shared genetic drift between the study populations from Southwest India and several Eurasian populations using an outgroup *f_3_*-statistic of the form *f_3_(Study_populations, X; Mbuti)*, where *Study_populations* correspond to one of the Southwest Indian populations sequenced in this study and *X* to a set of Eurasian populations from the dataset described in Methods. We observed a higher genetic affinity between all our study populations and other Indian populations (Figure 2D, Figure S7A-E, S8A-E), particularly from South India, which is consistent with the results of PCA and ADMIXTURE. More specifically, patterns of shared drift between the study populations and other South Indian populations differed, as seen in previous analyses. The Kodava, Kodava_US, Bunt, and Nair shared higher genetic affinity with each other and, to an extent, with populations that derive genetic ancestry from both ANI- and ASI-related sources, while the Kapla were genetically similar to populations with more ASI-related genetic ancestry such as tribal populations like Ulladan from Kerala^70^ and individuals from Handigodu village of Shimoga District of Karnataka^14^.

We constructed a maximum-likelihood tree using Treemix and modeled up to ten admixture events using a subset of present-day and ancient populations from Eurasia and an African outgroup (Mbuti), excluding populations with known recent admixture from Africa and East Asia (Figures S9A-K, Table S5). Without migration edges, we observed expected broad-level relationships between the populations, with Western and Eastern Eurasian populations forming distinct clusters. South Asian populations fell between these clusters, with populations that maximized ANI-related genetic ancestry reflecting higher similarity to Western Eurasians and populations that maximized ASI-related genetic ancestry being more similar to the Onge and East Asians populations. The Kodava, Kodava_US, Bunt, and Nair were closely related to each other and to other South Indian populations such as Iyer and Iyangar. The Kapla clustered with populations with a higher proportion of ASI-related ancestry, such as Paniya and Palliyar, consistent with previous analyses (Figure 2B). Few early migration edges (Table S5) recapitulated previously-reported events such as the Steppe-related gene flow, represented by ancient Central Steppe MLBA individuals from Russia, into the French.

### Estimation of admixture proportions and timing in Southwestern India

To directly model the study populations as a mixture of genetic ancestral components, we used both the *f_4_* ratio test and qpAdm. A two-way admixture model tested using the *f_4_* ratio method has been shown to be overly simplified for most Indian populations ^15,21,30,71^, and we used it primarily to assess the relative proportions of Central Steppe MLBA related genetic ancestry, a proxy for ANI-related genetic ancestry, in the study populations. The proportion of Central Steppe MLBA in the Nair, Bunt, Kodava, and Kodava_US ranged between 45% (+/− 1.4%) and 48% (+/− 1.3%), similar to the proportion found in other populations with higher ANI-related genetic ancestry, including some South Indian and the majority of North Indian populations. In contrast, the proportion of Central Steppe MLBA in Kapla was genetically similar to neighboring South Indian tribal populations with higher ASI-related genetic ancestry (27.17% +/− 2.1%), which was lower than the other study populations (Figure S10).

We next investigated more complex admixture histories, particularly three-way admixture related to Central Steppe MLBA, Indus Periphery cline (primarily ancient Iranian farmer-related ancestry) and Onge. We employed the qpAdm framework starting from a previously proposed model by Narasimhan *et al*. ^15,38^. As observed in ^15^, several South Asian populations, including the two Kodava groups, Nair, and Bunt sequenced in this study, have p-values less than 0.05 ^38^ (Table S6). This suggests the tested model does not sufficiently explain the genetic ancestries in these four populations as well as in many other South Asians. We further included as additional sources various Scythian groups and proxies of groups contemporaneous to Alexander the Great’s reign, as indicated in the oral histories of these populations, but also failed to retrieve working models for these tests (Table S7).

In contrast, the Kapla could be modeled with little Central Steppe MLBA-related genetic ancestry (4.7%) compared to genetic ancestries related to Indus Periphery Cline (40.8%) and Onge (54.6%). These genetic ancestry proportions were similar to Palliyar, Paniya, and Ulladan, which are South Indian tribal groups with higher ASI-related genetic ancestry (Table S6). Furthermore, we found limited evidence for Siddi (or African) ancestry contributing to the Kapla, based on the above qpAdm results as well as admixture *f_3_*-statistics, with no *f_3_* statistics reflecting a significant genetic contribution from a Siddi-related or African genetic ancestry source (Table S8). We additionally explored the increased variation in PCA among the Kapla by separating Kapla individuals into two groups based on the PCA results where Kapla_A included 6 individuals closer to the ASI end of the cline and Kapla_B included the other 4 individuals closer to the ANI end. We estimated D-statistics of the form *D(Kapla_B, Kapla_A; H3, Mbuti)*, where H3 denotes populations from Europe, the Middle East, and populations from South Asia that fall on the ANI-ASI cline. In agreement with the PCA, we found that most tested European and Middle Eastern populations were significantly closer to the four Kapla individuals closer to the ANI end (Kapla_B), and a few tested populations that maximized ASI ancestry, such as Ulladan, Paniya, and Palliyar, were significantly closer to Kapla_A (Table S9). To infer the proportions of Central Steppe MLBA, Indus Periphery Cline and Onge in Kapla_A and Kapla_B, we implemented qpAdm using sources and outgroups from ^15^ (Table S6). Kapla_A exhibited a slightly higher proportion of Onge-related ancestry (57.7%) compared to Kapla_B (49.8), and Kapla_B exhibited a slightly higher proportion of the other two components: 7.6% Central Steppe MLBA versus 2.8% in Kapla_A and 42.6% Indus Periphery Cline versus 39.5% in Kapla_A. We could not further narrow down the mode or timing of the within-population differentiation we observed above (e.g., via recent admixture with an ANI group or more ancient structure) due to the small sample size and substantial ancestry sharing between Kapla and neighboring South Indian populations included in our tests.

Broadly, the genetic source proportions in the study populations were within the range displayed by other South Asian populations, though the genetic ancestries related to Central Steppe MLBA and Indus Periphery were qualitatively at the higher end of the range in the Nair and Kodava compared to those observed in neighboring Indian populations. The relative amounts of allele sharing of our study and other South Asian populations with the three genetic sources, determined using *D*-statistics (Figure S11A-C, S12A-C), were in agreement with these results and our other analyses. More genetic data, especially from ancient samples from the region, can potentially shed further light on the complexities in genetic ancestry that may be missing in the current model.

Finally, to estimate the timing of the introduction of genetic ancestral source(s) related to ANI in South Asia, we ran ALDER on select South Asian populations using PC-derived weights reflective of the ANI-ASI ancestry cline to anchor the ancestry sources ^15,21,45^. The date estimates for populations with higher Central Steppe MLBA-related genetic ancestry, typically from North India, ranged from 60.4 ± 5.82 to 116.8 ± 13.37 generations. South Indian populations with higher ASI-related genetic ancestry tended to display slightly older dates than the former group, ranging from 92.34 ± 6.64 to 196.36 ± 48.6 generations (Table S10), in agreement with previous studies ^15,21^. Excluding the Kapla that did not yield a significant result (Z-score >=2, corresponding to a p-value < 0.05), the ANI-ASI admixture time estimates for the populations sequenced in this study ranged between 94.79 ± 6.68 and 111.23 ± 9.29 generations, which was within the range displayed by other populations included in the analysis^45^ (Table S10). As pointed out previously, these dates represent a complex series of admixture events and, particularly for populations with both the Indus Periphery Cline and Central Steppe MLBA-related genetic ancestries, may reflect an average of these events ^15,21^.

### Characterizing fine-scale genetic admixture using haplotype-based analyses

To evaluate population structure and genetic affinities at a finer scale, we implemented a haplotype-based analysis using ChromoPainter and fineSTRUCTURE ^47^. In this analysis, we first investigated general patterns of haplotype similarities followed by regional level patterns, including only South Indian populations. The degree to which an individual copies their haplotypes from another individual reflects the genetic similarity between those individuals at a haplotypic level.

At a broad level, we observed that both the Kodava groups, Bunt, Nair, and Kapla had the highest haplotype-copying rates from South Asian ancestry donors, particularly South Indian populations, consistent with previous analyses (Figure S13). At a regional scale, the populations sequenced in this study clustered into three groups: i) Nair and Bunt, ii) Kodava, Kodava_US, and Coorghi, and iii) Kapla (Figure 3 and Figure S14). Within the first two clusters, we could not identify further discernable sub-structure. When evaluating haplotype similarity to Central and Western Eurasian sources of ancestry, we found that the Nair & Bunt have a qualitatively higher per-locus copying probability to these sources relative to the Kodava, though not quantitatively significant (Figure S15, Table S11). In the case of Kapla, we observed haplotype similarity with Ulladan and Handigodu village donors, and significantly lower per-locus copying to Eurasian ancestry sources (Figure S14). The additional sub-structure uncovered through haplotype-based methods across the Nair, Bunt, and Kodava groups suggests more recent population structure in these populations.

**Figure 3:**
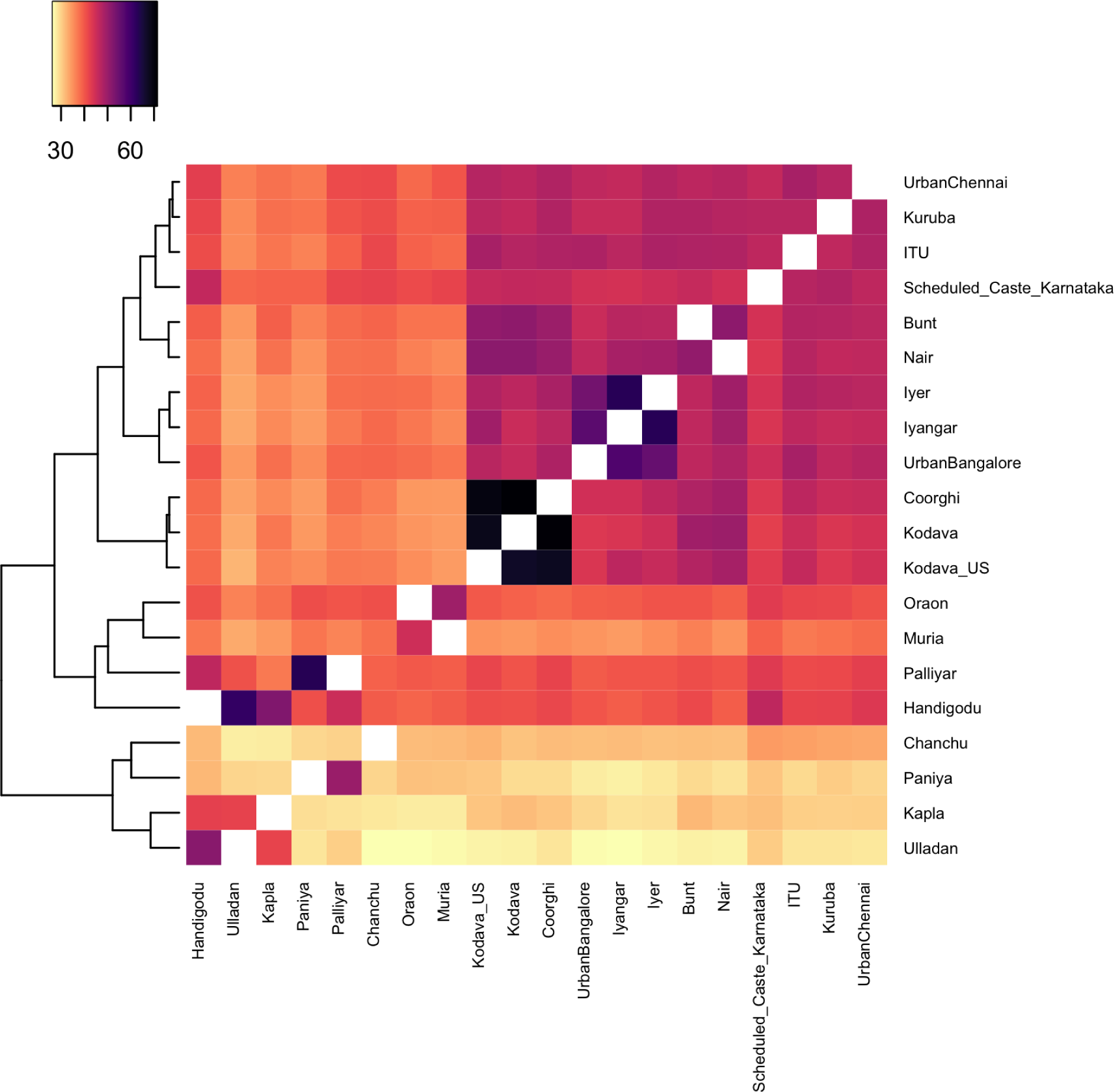
Haplotype-copying co-ancestry matrix across the regional level representing South Indian populations. Color bar represents the average length of haplotypes in centiMorgans (cM) shared between populations.

### Uniparental marker diversity in the Southwest Indian populations

The populations sequenced in this study displayed a mix of maternal lineages found today in South Asia (haplogroup M) and Western Eurasia (haplogroups R, J, U, HV) (Figure 4A, Table S12). Mitochondrial DNA haplogroup (mtDNA hg) R, reported to have originated in South Asia ^72–75^, was observed in each of the study populations. Individuals from both Kodava groups, Bunt, and Nair carried lineages of mtDNA hg U, previously reported in ancient and present-day individuals from the Near East, South Asia, Central Asia, and Southeast Asia ^15,76,77^. Some of the newly sequenced individuals belonged to subclades of haplogroup HV, a major subclade of haplogroup R0 found broadly from Eastern Europe to South Asia ^15,78,79^. Additionally, two Kodava_US individuals were assigned to mtDNA lineages that have, to our knowledge, not been observed in South Asia thus far. These included a subclade of haplogroup J (haplogroup J1c1b1a) reported in ancient individuals in Central and Eastern Europe ^71,80^, and haplogroup A1a, a subclade of haplogroup A primarily observed in East Asia ^81^.

**Figure 4:**
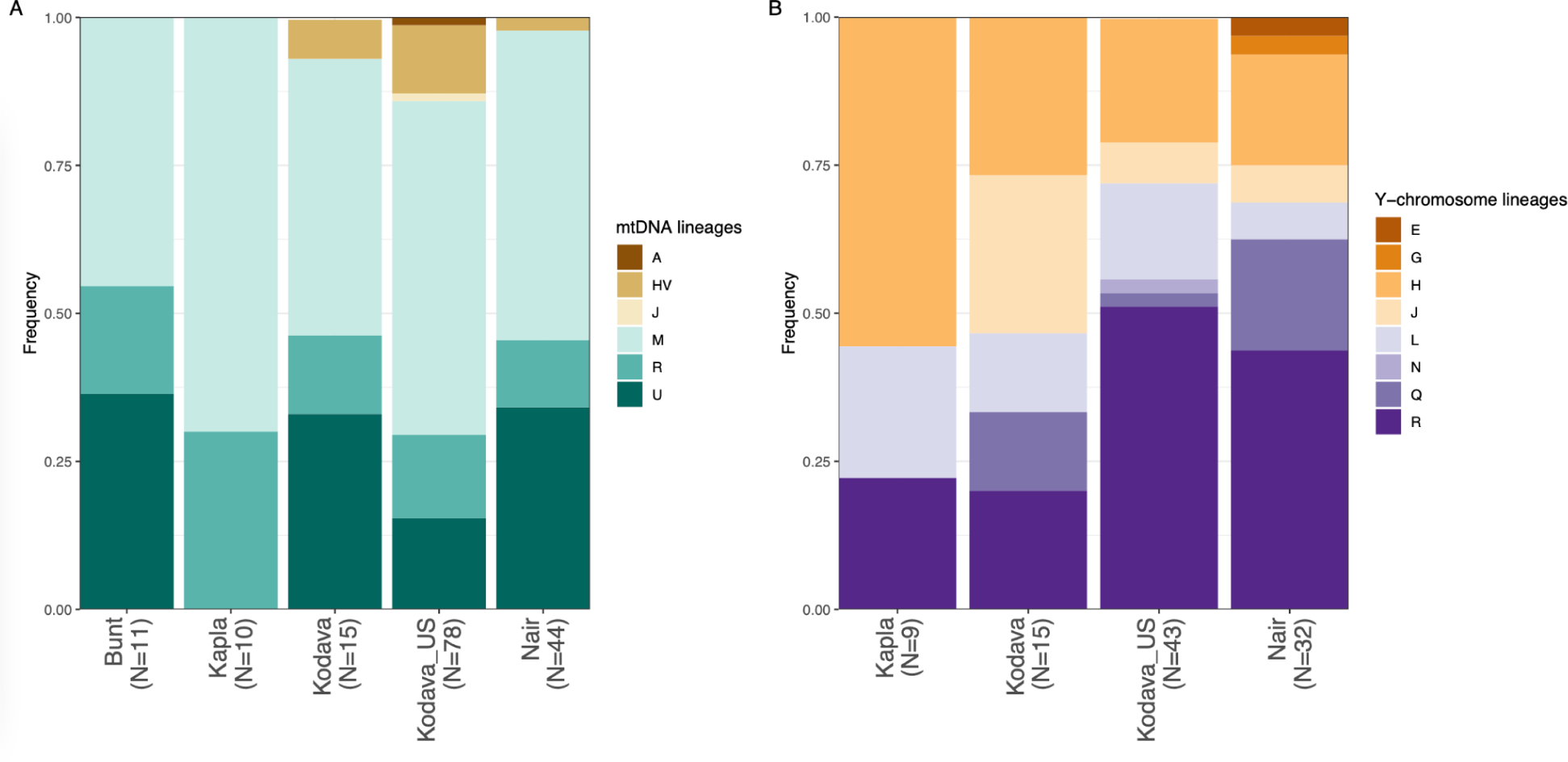
Distribution of the major mtDNA **(A)** and Y chromosome **(B)** haplogroups from the analysis of 158 complete mitochondrial genomes and 99 complete Y chromosomes from this study. Bunt were not included in the Y chromosome analysis due to a single male in the dataset.

We observed a total of eight Eurasian Y-chromosome haplogroups across 99 newly sequenced male individuals (R, H, L, G, N, J, E, Q) (Figure 4B, Table S13). R, H, and L Y-chromosomal lineages have been previously reported in present-day and ancient individuals from South Asia, Southeast Asia, Central Asia, the Arabian Peninsula, Europe, and present-day Turkey ^15,82–86^, while haplogroups G, N, J, E, and Q are found more broadly across Eurasia ^87,88^.

To explore the relationship between uniparental markers and matrilocality for the Nair, we compared the haplotype and pairwise diversity on the mitochondrial DNA (mtDNA) similar to analyses from ^52^ (Table S14). For groups with similar sample sizes, we found the strongest difference in haplotype diversity to be between the Bunt and Kodava samples (p = 0.19; t = −0.88), which does not reject the null hypothesis of similar haplotype diversity matrilocal and patrilocal groups in our dataset.

### Endogamy in Southwest Indian populations

Founder effects and endogamy are prominent demographic forces shaping genetic diversity in populations of South Asian genetic ancestries ^14,20,89^. We sought to characterize rates of endogamy in the study populations relative to other populations sampled in India as endogamy can increase the frequency of recessive disease alleles. We calculated the identity-by-descent (IBD) score, which measures the extent to which unrelated individuals in a population share segments of the genome identical-by-descent (Figure 5).

**Figure 5:**
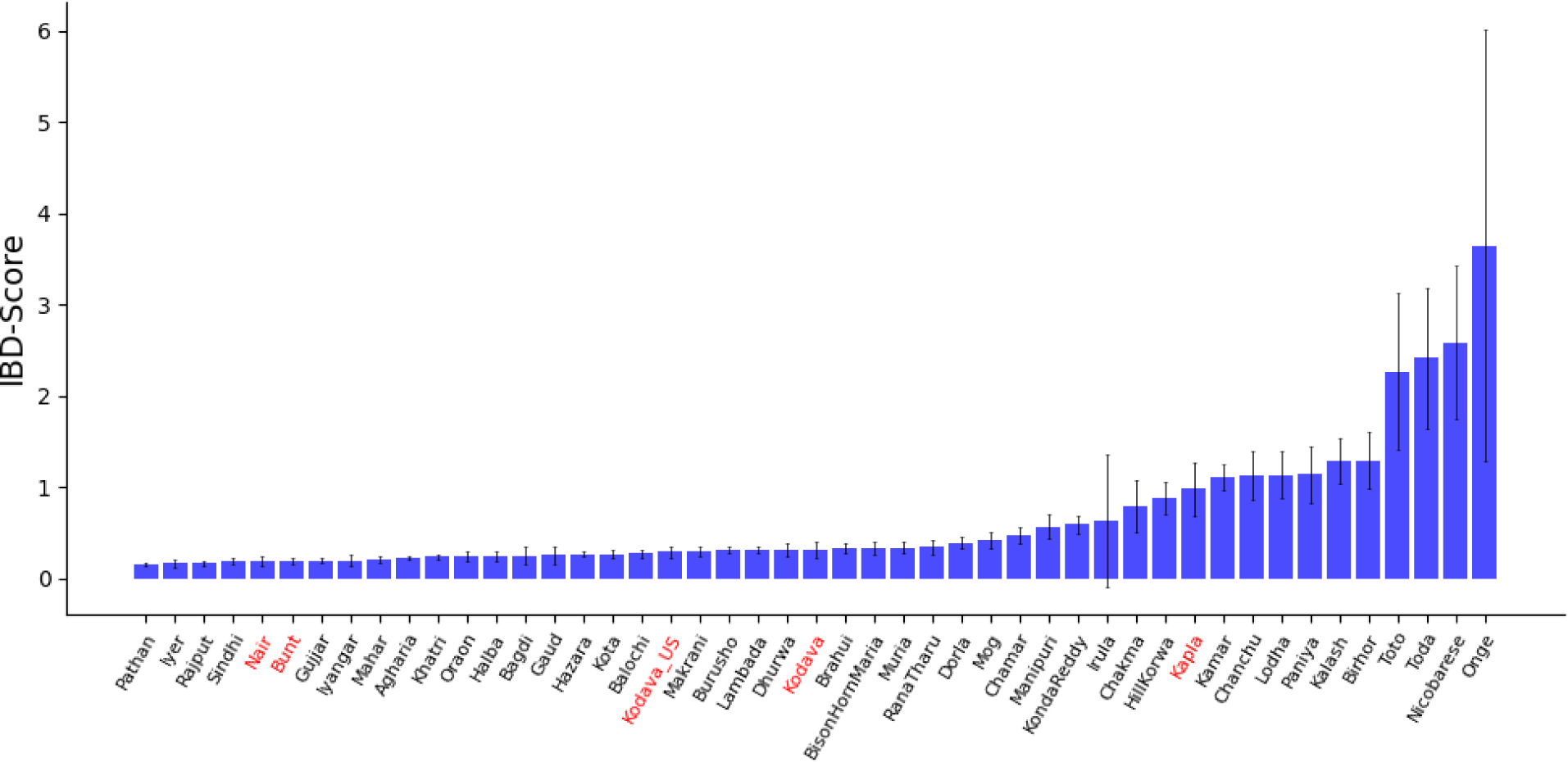
IBD score across multiple populations in South Asia. Populations sequenced in this study are highlighted in red.

We found that the Nair and Bunt populations had similar levels of endogamy as captured by the IBD score. Additionally, both Kodava populations had higher IBD scores than the Bunt and Nair but were consistent between themselves. We found the Kapla to have elevated levels of endogamy relative to geographically neighboring populations such as the Kodava, but similar to several tribal groups from Central and South India.

We estimated runs of homozygosity (ROH) to further explore endogamy in the study populations. In agreement with the IBD scores, we observed the Kapla had an increased number of ROH (NROH) and generally longer ROH (>10Mb) (Figures S16 and S17), consistent with smaller effective population size and higher levels of consanguinity ^57^. To corroborate these results, we also estimated the occurrence and magnitude of founder events in the newly sampled populations using ASCEND ^55^. We detected a recent founder event for the Kapla occurring ∼10 generations ago and with a founder intensity of ∼12%, but the Nair, Bunt, and both Kodava populations did not show signs of recent founder events (Figures S18). In summary, our results suggest modest levels of endogamy within our newly sampled populations, with the Kapla reflecting a novel founder effect.

### Novel variant discovery in Southwest Indian populations

We generated high-coverage genomes (>30x) for a total of nine individuals representing the Bunt, Kapla, Kodava_US, and Nair, and evaluated the presence of variants not previously categorized in existing catalogs of genetic diversity. After alignment, quality control filters, and genotype calling across these WGS data, we ran Ensembl VEP v105.0 to assess whether each variant was found in pre-existing catalogs of variation and their putative functional consequences. We used 1000 Genomes, GnomAD, and GenomeAsia as catalogs of allelic variation against which to compare and establish novel occurrences of variants. We observed 11,132,650 total variants (8,562,627 SNPs, 2,570,023 indels) and, of these, 208,041 (151,137 SNPs, 56,904 indels) were not found in existing catalogs of variation, which we term as “novel variants”. Of these novel variants, we assessed the proportion of SNPs and indels in specific functional impact classes as defined by VEP (modifier, low, medium, high). For novel SNPs, there were 150,315 modifier, 349 low, 444 moderate, and 29 high impact variants. For novel indels, the proportions were similar with 56,714 modifier, 49 low, 36 moderate, and 105 high impact variants. We also observed that the majority of the putatively high impact variants were singletons, i.e. only found on a single haplotype out of the 18 that are tested, which is to be expected based on the action of negative selection ^90^.

We also explored the burden of moderate and high effect variants within this small set of high-coverage genomes, given previous evidence of endogamy in Indian populations influencing the burden of deleterious mutations ^14,19^. When looking at only observed mutations with moderate or high predicted effects based on VEP, we found that there was an enrichment of deleterious variants in the Kodava_US population (p < 1e-12; Mann-Whitney U: 24e6 between Kodava_US and Bunt), which does not support a direct relationship between predicted endogamy (Figure 5) and burden of deleterious variants (Figure S19A). However, this is confounded by the fact that the Kodava_US samples had ∼77x coverage compared to 30x coverage from the other populations, so there was a higher absolute number of variants discovered in the Kodava_US individuals.

We hypothesized that populations with higher fractions of ASI ancestry would have a higher fraction of novel variants due to undersampling of ASI-related genetic ancestry in known catalogs of human allelic variation. We found a strong enrichment of novel variants in the Kapla, supporting our hypothesis (p < 1e-12; Mann-Whitney U: 25e6 between Kapla and Bunt). In absolute terms, this is a difference of ∼ 3,000 variants per haplotype (Figure S19B). Overall, these findings motivate conducting further whole-genome surveys of genomic variation in underrepresented genetic ancestries both for discovery of functional genetic variation and improving catalogs of allelic diversity ^e.g. 19^.

## Discussion

This study engaged with the genetic and oral histories of populations from Southwest India. By analyzing whole-genome sequences from participants in India and the US, we reconstructed broader and fine-scale genetic relationships of the study populations to worldwide and other South Indian populations, respectively. Surveys of population oral histories and origin stories were conducted through community interactions, augmented by published historical and limited anthropological works. In the context of this study, we did not differentiate between oral history and oral tradition as many anthropologists do, and instead used oral history as an umbrella term to encompass aspects of cultural histories of the study populations that have both been passed down orally from one generation to the next and drawn from contemporary experiences and observations. Furthermore, oral histories and self-identities are inherently complex and may be shaped over time by community and external influences. Hence, we focused on those aspects of oral histories that specifically relate to origins and connections with other communities as recognized by community members today.

We found the Bunt, Kodava, and Nair to share close genetic ancestry with each other and with other neighboring South Indian populations. These populations, along with several other Indian populations, have genetic ancestries that can be modeled with Onge, Indus Periphery Cline, and Central Steppe MLBA populations. This mix of genetic ancestries is reflected in the uniparental markers that also show a mix of South Asian and Western Eurasian mtDNA and Y chromosome haplogroups. Future studies are needed to better understand the distribution ranges of haplogroups not typical of this region, such as mitochondrial haplogroups A1a and J1c1b1a.

Our result of not detecting additional sources of Central and/or Western Eurasian ancestry in the Bunt, Kodava, and Nair motivates anthropological follow-up to better understand the connections in their oral histories to non-local populations such as ancient Scythians and members of Alexander’s army. It is possible that our sampling scheme and/or analytical power limits the detection of such a signal in the genetic data, particularly if subsequent admixture with local Indian populations caused a dilution of non-local genetic signatures in these populations. Moreover, the timing of ANI-ASI admixture, reported to be between 2,000-4,000 years before present ^21^, may confound detection of admixture events noted in oral histories, such as that between local Indian populations and members of Alexander’s army sometime after 327 BCE. In fact, the Nair and Kodava individuals sequenced in this study have a slightly higher proportion of Western Eurasian ancestry compared to neighboring populations in South India, though comparable to and/or lower than most North Indian populations. The available data in this study precludes determination of the reason for this slightly elevated signal in these populations, including potential sampling bias and long-standing endogamy.

The observed discordance between oral and genetic histories leads to interesting avenues for ethnographic and anthropological follow-up while, at the same time, providing a valuable opportunity to consider the dangers of conflating self-identities with genetic ancestries. Could these origin stories be reflective of past cultural and/or economic contacts rather than involving gene flow? In studies of demographic histories, the expectation of origin stories arising from oral traditions to converge with genetic histories, by using the former to motivate hypotheses that are then either accepted or rejected by the genetic data, tends to suggest that genetic and oral histories should be alignable. On the contrary, they represent distinct facets of individual and group identities that should not be forced to reconcile by pitting one against another ^10,12^ or be assumed to occur across similar timeframes. Taking a similar stance, we stress that our findings based on genetic data should not be used to validate or reject community origin stories based on oral traditions. Instead, we encourage further anthropological research on the nature of the relationships with non-local populations prevalent in the oral histories of the populations included in our study. This would serve to enhance our understanding of the cultural and oral legacy of these populations and potentially illuminate new dimensions of the regional socio-cultural history. Such conversations are especially pertinent today as the genomics revolution coupled with direct-to-consumer initiatives over the past two decades has brought conversations around genetics and ancestry squarely into the public domain and calls for increased engagement from researchers to explain the interpretive scope of the data ^91^.

At a more fine-scale level, haplotype-based analysis suggests more recent genetic contacts between Bunt and Nair populations. Moreover, the close genetic relationship observed between the Kodava from India and Kodava_US supports these individuals deriving from the same Kodava population in India. Although we lack socio-cultural context for the Coorghi individuals genotyped in ^14^, we conjecture that the genetic similarity of the Coorghi and the two Kodava datasets from this study may, again, be capturing their common origin since Coorghi is an anglicized version of Kodava that was adopted during the colonial times. Overall, close genetic affinities between the Bunt, Kodava, and Nair do, in fact, complement their oral histories that speak to historical cultural contacts as well as overlapping self-identities, likely facilitated by their geographical proximity to one another. Our genetic analyses do not detect any discernible sub-structure within the Bunt, Kodava, and Nair despite the Nair and Kodava donors originating in different locations within Kerala and Kodagu, respectively, and the complex socio-cultural subgroups recognized within the Nair population. This may suggest that geography and social stratification have not produced long-term reductions to gene flow within these populations.

We also found little statistical support from genetic diversity statistics (Table S14) to distinguish between groups anthropologically associated to be matrilocal. From haplotype and nucleotide diversity on mtDNA, we did not find statistically significant lower levels of these metrics expected for matrilocal groups, especially the Nair, compared to patrilocal groups. From haplogroup diversity on the Y chromosome relative to the mtDNA, the Nair display a slightly higher diversity than the Kodava, which may be expected under matrilocality. However, this result is qualitative and it is likely that larger sample sizes per group would be required in order to address this in a stronger quantitative sense. There are other possible explanations for a lack of clear signal in uniparental markers between matrilocal and patrilocal populations, such as recent shifts from strict matrilocality that could potentially dilute the signal in the mtDNA genetic diversity, or smaller long-term population size that could result in shifts in diversity which may not be well-accounted for in our analyses. We leave these as future areas of work bridging matrilocal and patrilocal population dynamics to observed genetic data.

In contrast to the other study populations, the Kapla are genetically more similar to tribal South Indian populations, which have higher proportions of Onge-related genetic ancestry and negligible amounts of the Central Steppe MLBA genetic component. In this regard, the Kapla may represent one of several close genetic descendants of the ASI group ^15^. A close link of the Kapla to the ASI group is also evident in the uniparental markers, which are enriched for haplogroups with proposed South Asian origins. The lack of substantial support for a Siddi or African origin for the Kapla in the genetic data raises an interesting follow up to these links proposed in historical records. Furthermore, despite their geographical proximity to the Kodava and socio-economic relationships between these two populations observed by the research team, stark differences in lifestyle between the two populations may have resulted in more long-term genetic isolation of the Kapla from the Kodava. Their genetic isolation and elevated IBD score may be reflected in historical narratives, both published accounts that state the Kapla “*consist of only 15 families*” ^68^ and were relocated to their present location by a local ruler, and anecdotal accounts that suggest they were then isolated from neighboring populations.

However, we do detect slightly higher genetic ancestry related to Central Steppe MLBA and Indus Periphery Cline in four Kapla individuals relative to the others, which may be due to either recent contacts with neighboring populations, driven by socio-economic contacts between these populations, or latent population structure. Future studies with increased sample sizes are needed to further characterize the genetic variability within the Kapla more accurately.

Although our analyses were not designed to make claims regarding complex traits or disease, we expect to find an increased prevalence of recessive genetic disorders in populations with higher rates of endogamy ^14,20,92^. While follow-up studies with detailed phenotype information across populations in Southwest India will bring more evidence to bear on the relationship between endogamy and disease risk, particularly for recessive diseases, we found all four study populations to be within the range of IBD scores and ROH displayed by other Indian populations. Notably, the Kapla registered a higher IBD score and increased number and length of ROH compared to the Bunt, Kodava, and Nair, which suggests elevated endogamy in the Kapla, presumably due to their isolation and smaller population size. Our analysis of deleterious variants in a smaller, but higher-coverage, whole-genome dataset found that the Kapla were not the population with the highest burden of deleterious mutations. However, the Kapla did contain significantly higher rates of novel variants not found in existing public catalogs of allelic variation.

The results from this study provide an impetus for follow-up studies to further characterize the levels of genetic variation in these and other populations in India through increased sample sizes and phenotype collection. Additionally, future studies of population histories should engage closely with relevant fields such as anthropology. Importantly, such work should be conducted with substantial community engagement to ensure accurate recording of information on those aspects of their history that the population would like to further follow up on, given the sensitivities around identities stemming from oral histories.

## Supplemental Information

Supplemental Information includes 19 figures and 14 tables.

## Acknowledgments

First and foremost, we would like to thank members of the Kodava (both in India and in the US), Bunt, Nair, and Kapla populations for their participation in this study and for graciously hosting research team members in their communities and sharing their oral histories. We also thank members of the sequencing facilities at MedGenome Inc, Bangalore (India), Novogene, Sacramento (USA), and the University of Chicago DNA Sequencing Facility, Chicago (USA). We are also extremely grateful to Anna Di Rienzo, John Novembre, Matthias Steinrücken, and Bridget Chak for their valuable feedback on the manuscript and to Kendra Kodira for creating awareness and promoting participation among the Kodava_US community. This project was funded through the NIH Grant R35GM143094, University of Chicago start-up funds, Cincinnati Children’s Endowment, and the Gibbs Travelling Research Fellowship from the Newnham College at the University of Cambridge.

## Data availability

In compliance with the informed consent obtained from participants in the study, raw data (fastq files), alignments (bam files), and variant calls (VCF files) are available under data access agreement, requests for which should be submitted to: M.R. and N.R.

