## Supplementary Figures and Tables for "Distinct positions of genetic and oral histories: Perspectives from India"

| | | Page Number |
|---|---|---|
| Fig. S1 | $f_4$-ratio model. | 1 |
| Fig. S2 | *D*-statistics for $f_4$-ratio test. | 2 |
| Fig. S3 | PCA with pseudo-haploid calls. | 3 |
| Fig. S4A | Extended ADMIXTURE results across K=6 to K=11. | 4 |
| Fig. S4B | Cross-validation error of ADMIXTURE from K=6 to K=11, minimized in K=7. | 5 |
| Fig. S5 | ADMIXTURE with pseudo-haploid calls | 6 |
| Fig. S6A | PCA investigating genetic variation in Kapla | 7 |
| Fig. S6B | PCA to investigate the genetic relationship between Kapla and Siddi populations. | 7 |
| Fig. S7A-E | Outgroup-$f_3$ of the form $f_3$(Target, X; Mbuti). | 8 |
| Fig. S8 | Outgroup-$f_3$ of the form $f_3$(Target, X; Mbuti) using pseudo-haploid calls. | 9 |
| Fig. S9A-K | Treemix results. | 10-13 |
| Fig. S10 | $f_4$-*ratio* results | 14 |
| Fig. S11 | D-statistic with source populations from qpAdm model | 15 |
| Fig. S12 | D-statistic with source populations from qpAdm model using pseudo-haploid calls. | 16 |
| Fig. S13 | Summary of co-ancestry matrix as average length count per population from ChromoPainter. | 17 |
| Fig. S14 | fineSTRUCTURE clustering dendrogram | 18 |
| Fig. S15 | Ancestry-specific haplotype copying across sampled populations from Southwest India. | 19 |
| Fig. S16 | Total length of ROH (SROH) versus the total number of ROH (NROH) | 20 |
| Fig. S17 | Short versus long ROH for select populations | 21 |
| Fig. S18 | Decay curves for the Southwest India populations (Kapla, Bunt, Kodava, Kodava US, and Nair) and other South Asian populations | 22 |
| Fig. S19A-B | Functional and novel variation in high-coverage WGS samples from Southwest India. | 23 |
| Table S2 | Populations included in Treemix analysis. | 24 |
| Table S4 | Populations used for calculating PCA-based SNP-loadings. | 25-26 |
| Table S10 | The ANI-ASI admixture timings in select populations from India inferred using ALDER (fit started at d > 0.30 cM). | 27 |
| Table S14 | Haplotype and nucleotide diversity on mtDNA across matrilocal and patrilocal groups in Southwest India. | 28 |

**Figure S1: $f_4$-ratio model.** Representation of the model used for estimating the relative proportions of ANI- and ASI-related genetic ancestries in the Southwest Indian populations.

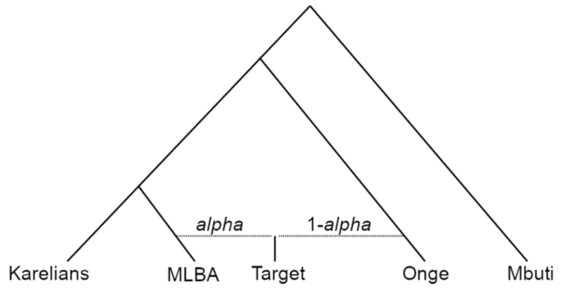

**Figure S2: *D*-statistics for *f₄*-ratio test.** *D*-statistics to evaluate the closest present-day population to Central Steppe MLBA (ANI proxy) for estimating α.

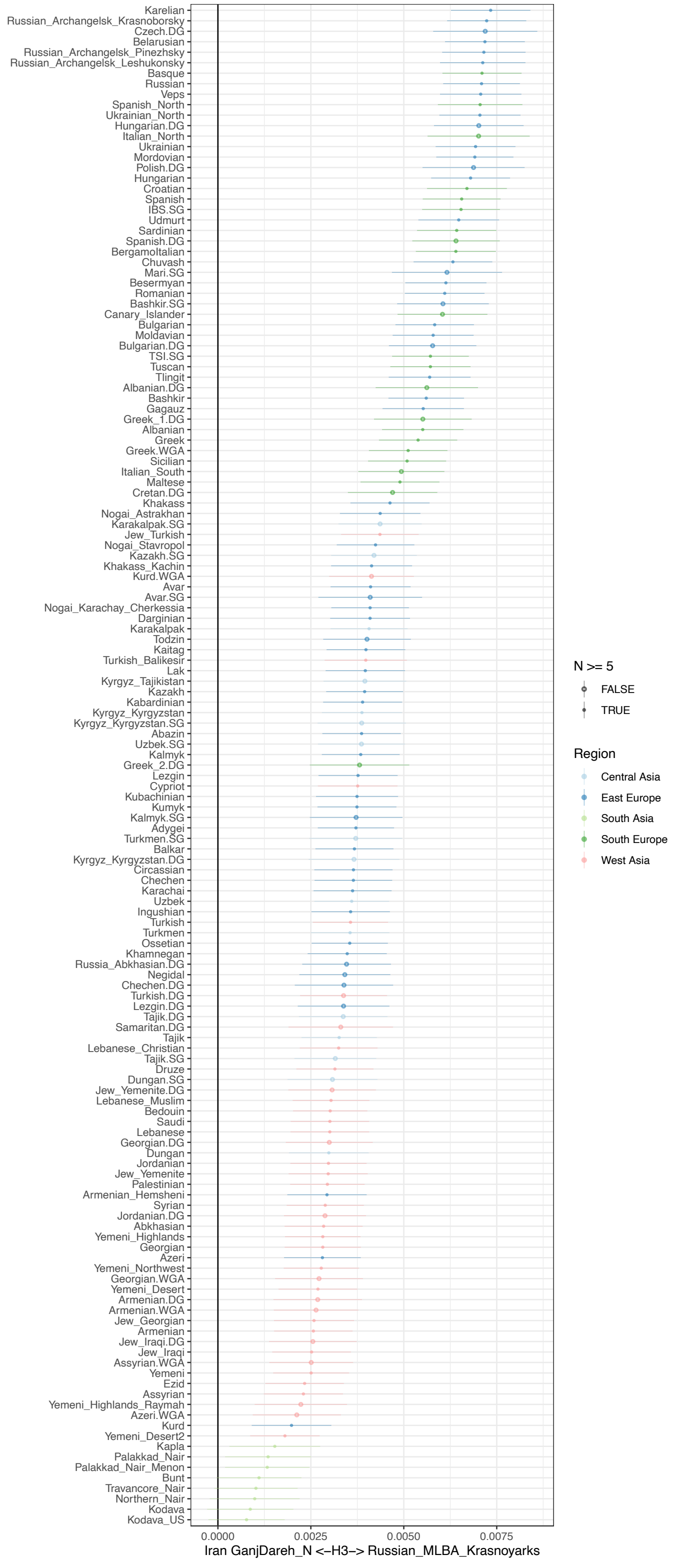

**Figure S3: PCA with pseudo-haploid calls.**

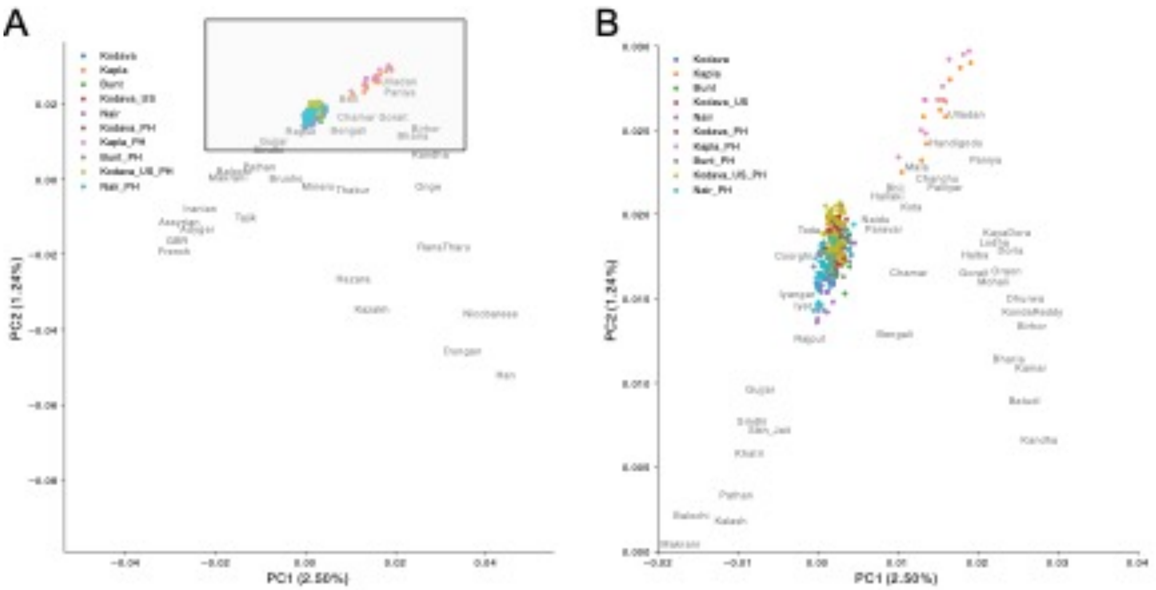

**Fig. S4A. Extended ADMIXTURE results across K=6 to K=11.**

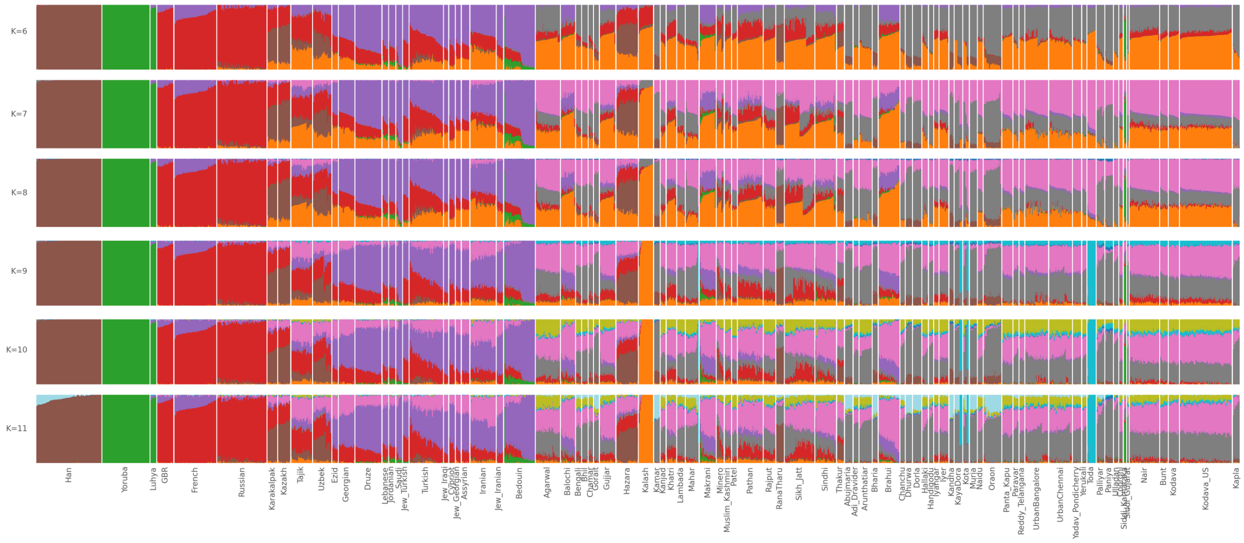

**Fig. S4B. Cross-validation error of ADMIXTURE from K=6 to K=11, minimized in K=7.**

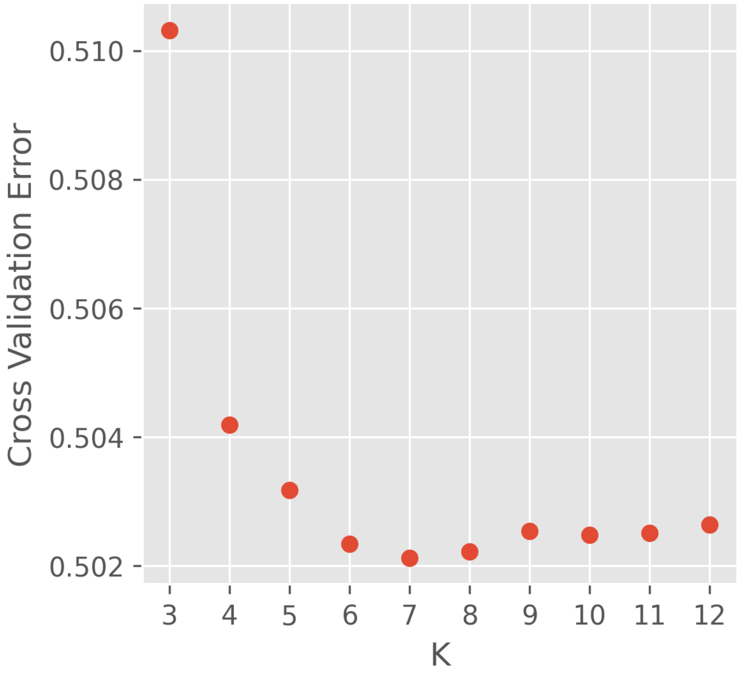

### Figure S5: ADMIXTURE with pseudo-haploid calls.

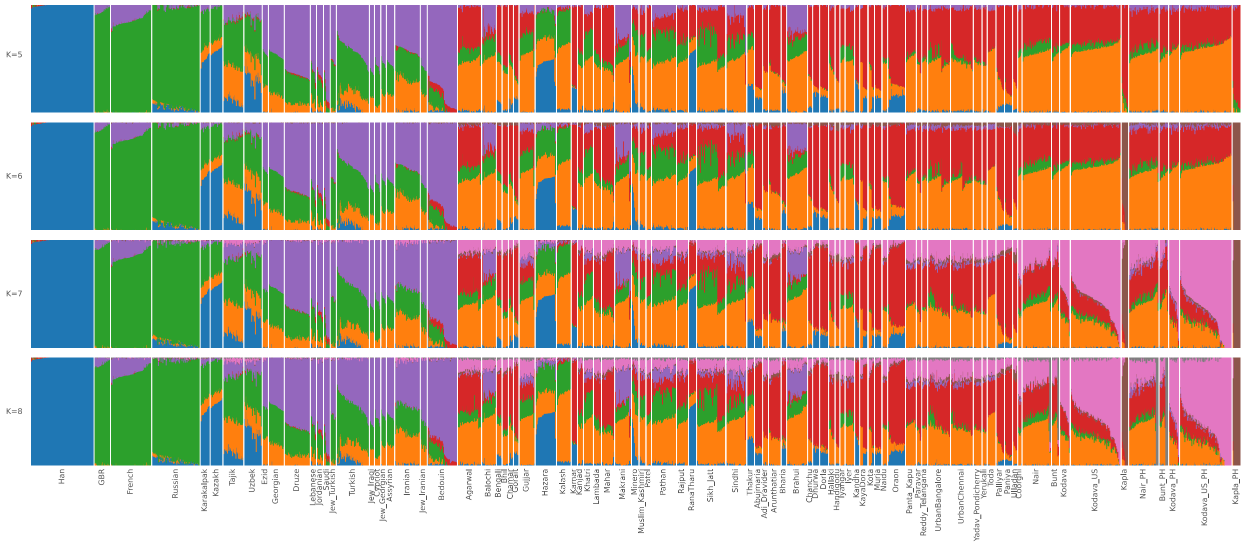

**Figure S6A: PCA investigating genetic variation in Kapla.** Dispersion of the ten Kapla individuals along the ANI-ASI cline with respect to other select South Indian populations with higher ASI component like the Ulladan, Paniya and Vysya. For comparison, Gujjar is shown as a representative of a population with higher ANI component.

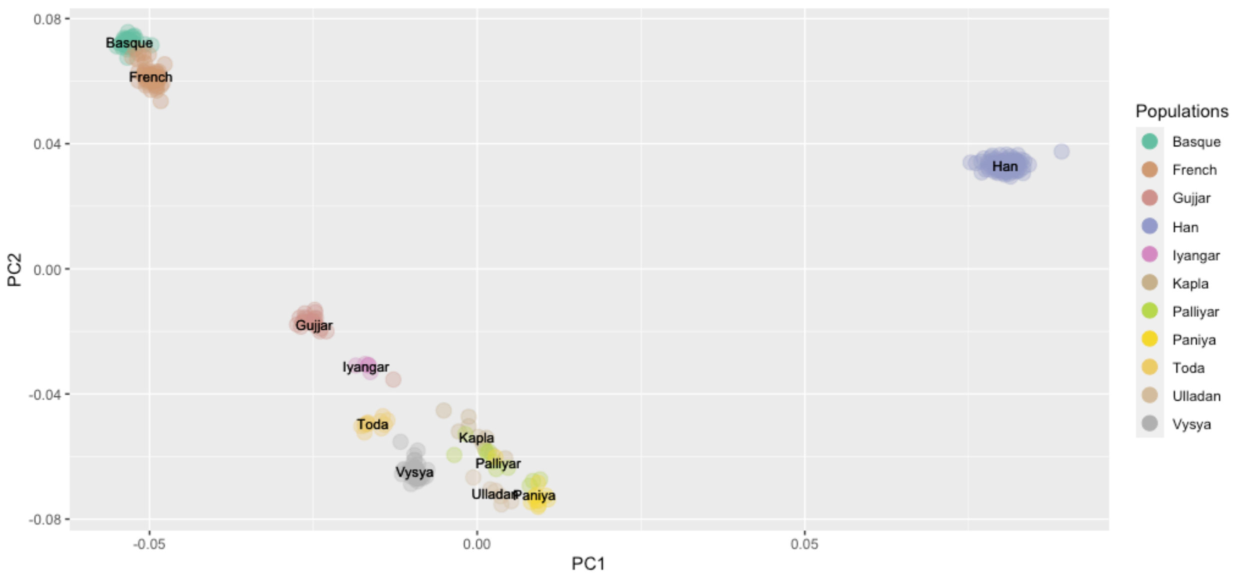


**Fig. S6B. PCA to investigate the genetic relationship between Kapla and Siddi populations.**

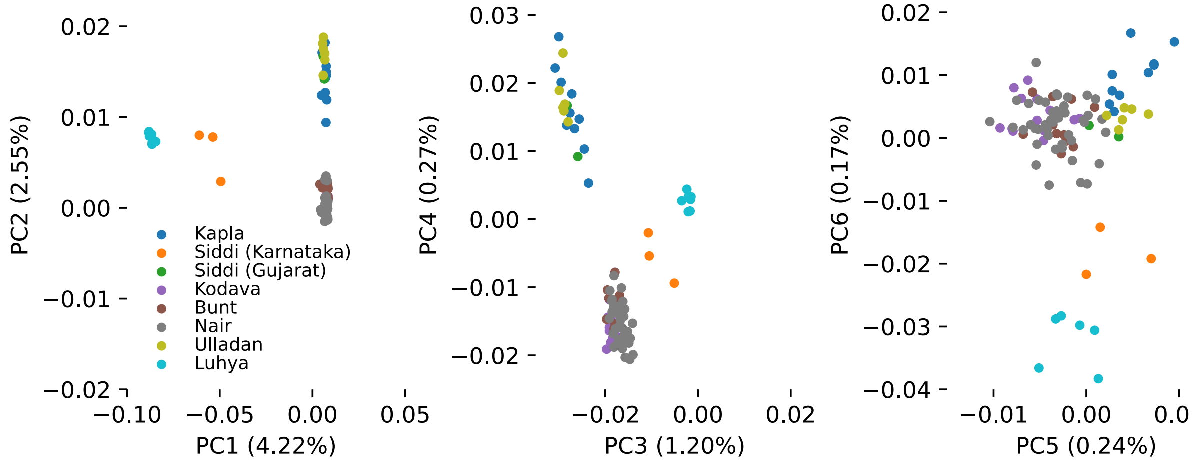

**Fig. S7A-E.**

**Outgroup-$f_3$ of the form $f_3$(Target, X; Mbuti).** Target corresponds to one of the Southwest Indian populations sequenced in this study and X to a set of Eurasian populations from the dataset described in Methods.

A

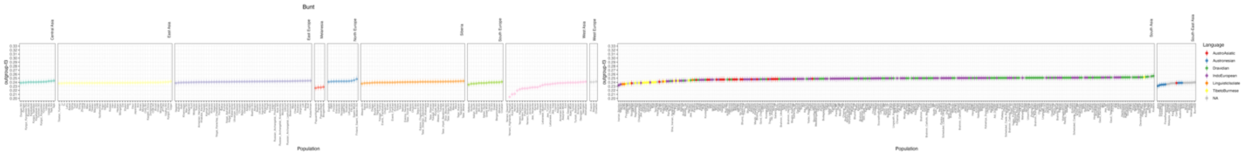


B

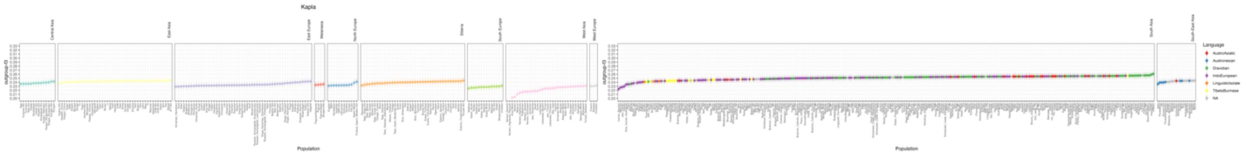


C

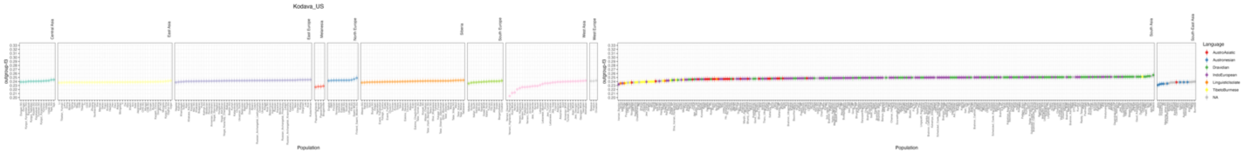


D

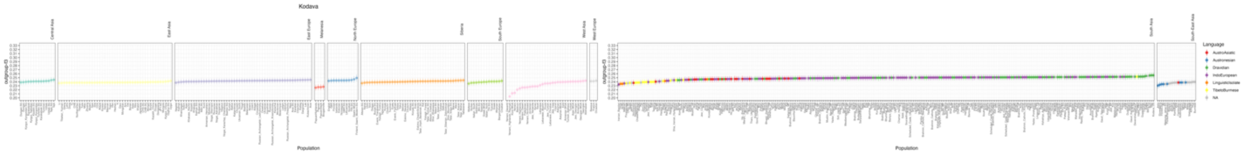


E

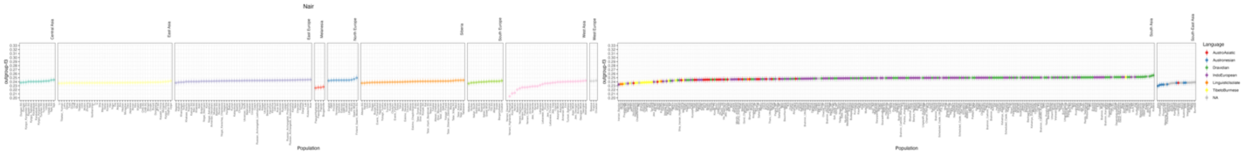

**Figure S8: Outgroup-$f_3$ of the form $f_3$(Target, X; Mbuti) using pseudo-haploid calls.** Target corresponds to one of the Southwest Indian populations sequenced in this study and X to a set of Eurasian populations from the dataset described in Methods.

**A)**

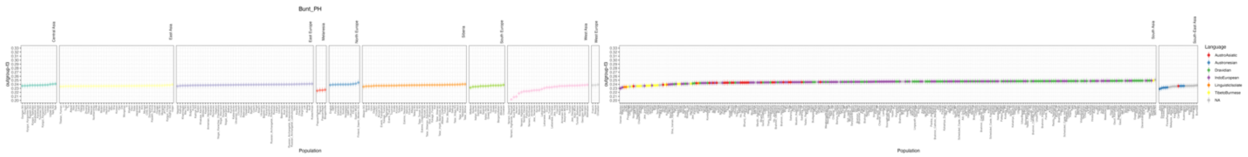


**B)**

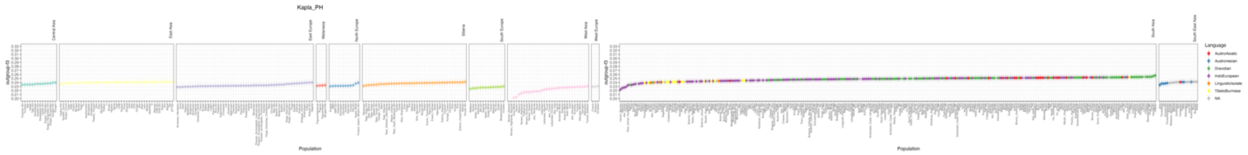


**C)**

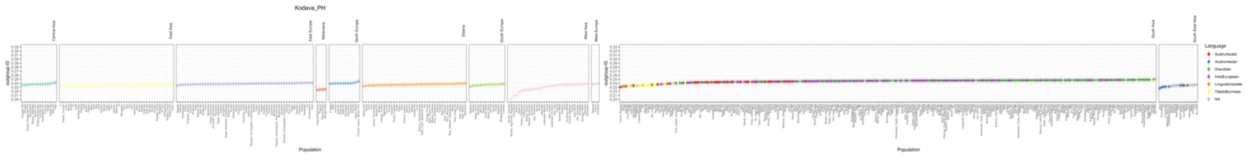


**D)**

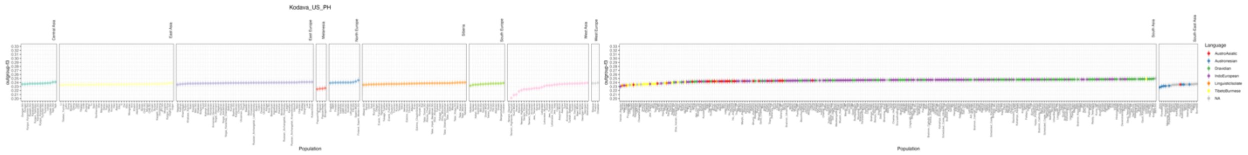


**E)**

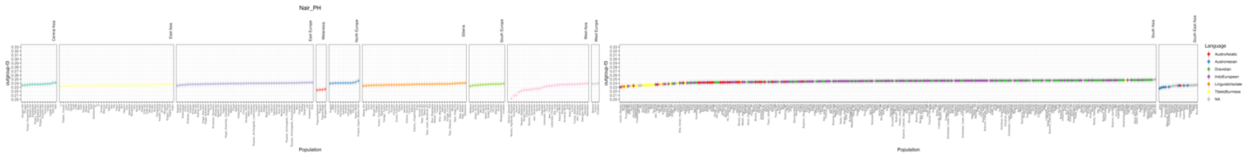

**Fig. S9A-K.**

**Treemix results.** Maximum Likelihood trees (left panel) and residuals (right panel) of allele frequency estimates from Treemix with up to 10 migration edges (m).

A) m=0

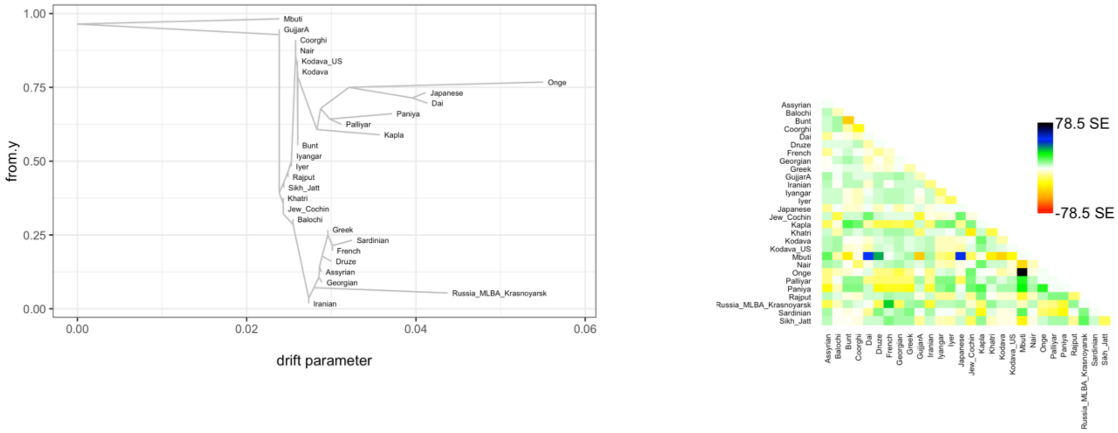


B) m=1

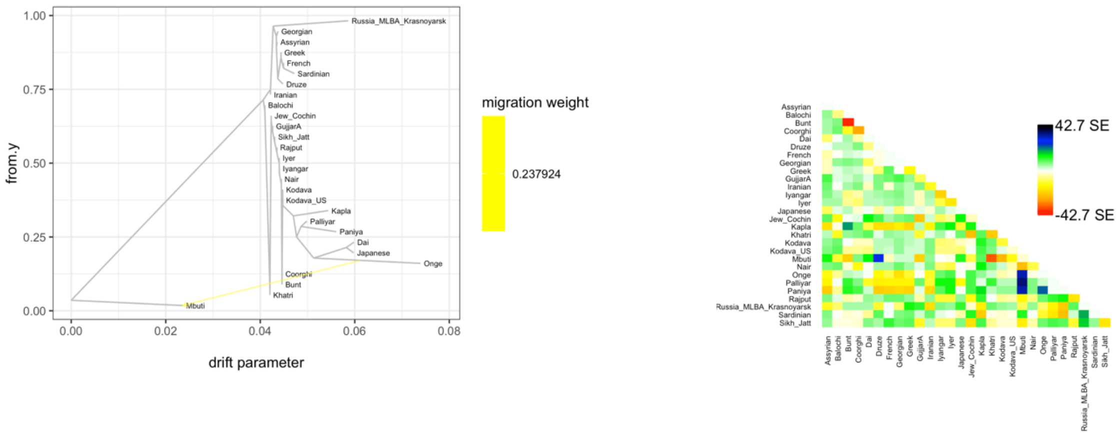


C) m=2

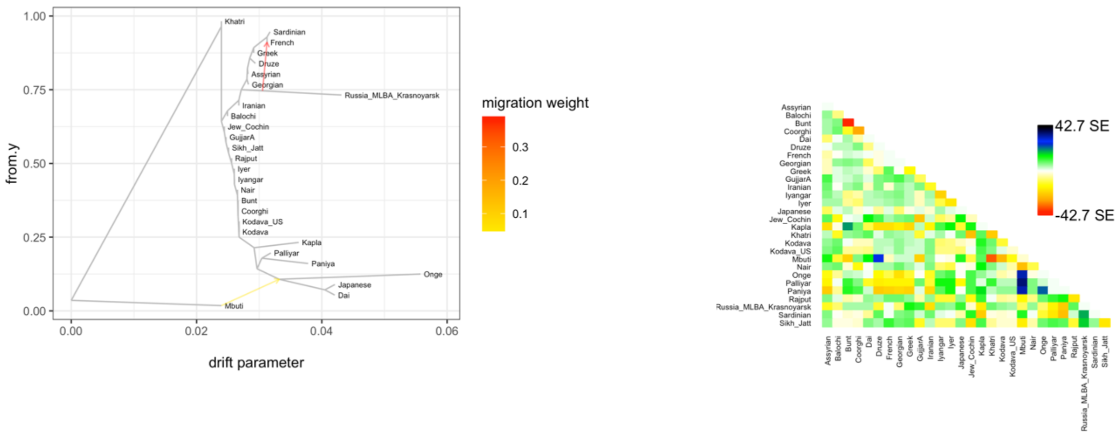

D) m=3

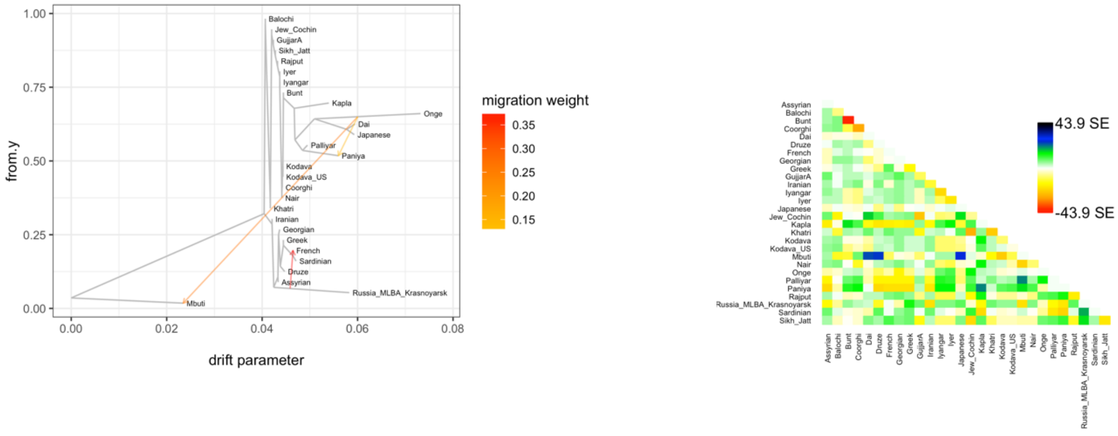

Balochi
Jew_Cochin
GujjarA
Sikh_Jatt
Rajput
Iyer
Iyangar
Bunt
Kapla
Onge
Dai
Japanese
Palliyar
Paniya
Kodava
Kodava_US
Coorghi
Nair
Khatri
Iranian
Georgian
Greek
French
Sardinian
Druze
Assyrian
Russia_MLBA_Krasnoyarsk
Mbuti
from.y
drift parameter
migration weight
0.35
0.30
0.25
0.20
0.15
43.9 SE
-43.9 SE


E) m=4

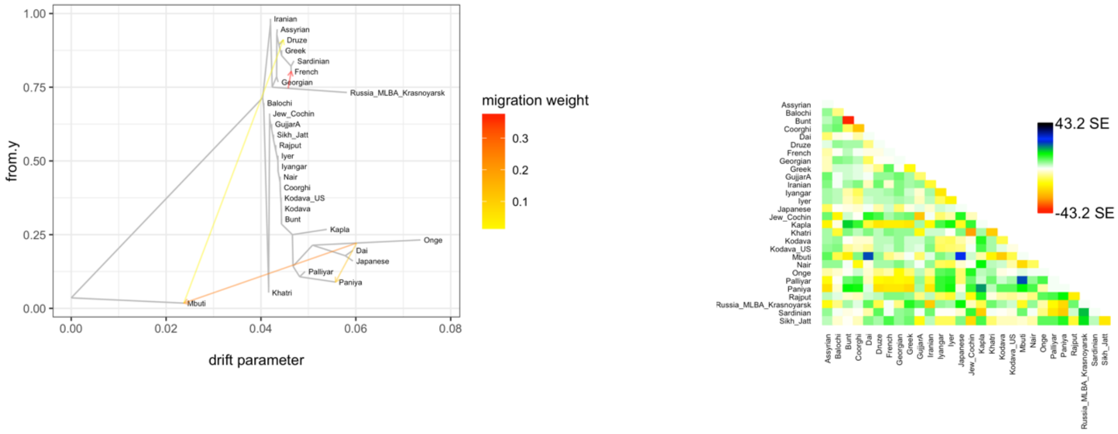

Iranian
Assyrian
Druze
Greek
Sardinian
French
Georgian
Russia_MLBA_Krasnoyarsk
Balochi
Jew_Cochin
GujjarA
Sikh_Jatt
Rajput
Iyer
Iyangar
Nair
Coorghi
Kodava_US
Kodava
Bunt
Kapla
Onge
Dai
Japanese
Palliyar
Paniya
Khatri
Mbuti
from.y
drift parameter
migration weight
0.3
0.2
0.1
43.2 SE
-43.2 SE


F) m=5

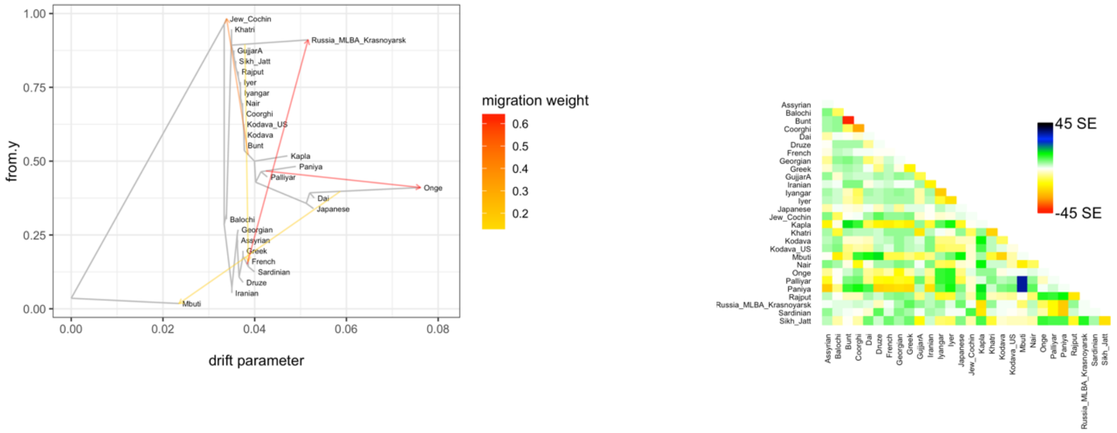

Jew_Cochin
Khatri
Russia_MLBA_Krasnoyarsk
GujjarA
Sikh_Jatt
Rajput
Iyer
Iyangar
Nair
Coorghi
Kodava_US
Kodava
Bunt
Kapla
Paniya
Palliyar
Onge
Dai
Japanese
Balochi
Georgian
Assyrian
Greek
French
Sardinian
Druze
Iranian
Mbuti
from.y
drift parameter
migration weight
0.6
0.5
0.4
0.3
0.2
45 SE
-45 SE

G) m=6

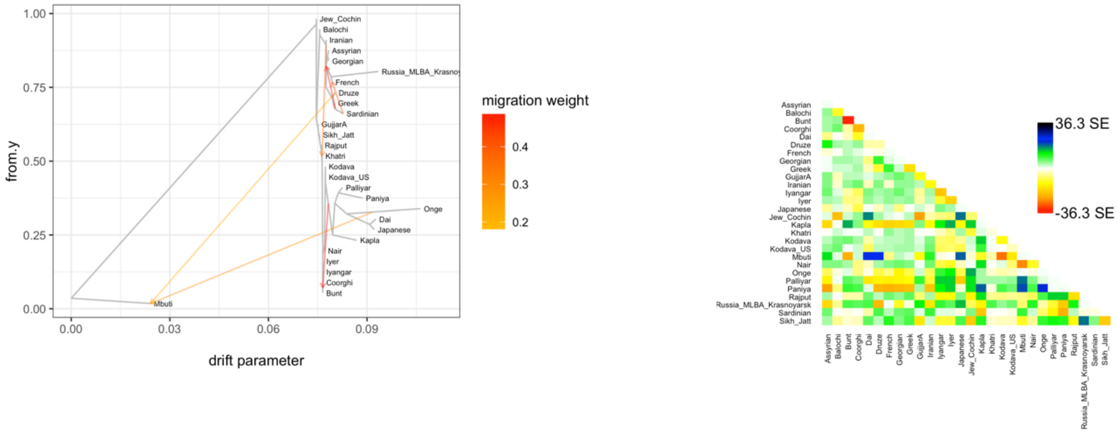
Jew_Cochin
Balochi
Iranian
Assyrian
Georgian
Russia_MLBA_Krasno
French
Druze
Greek
Sardinian
GujjarA
Sikh_Jatt
Rajput
Khatri
Kodava
Kodava_US
Palliyar
Paniya
Onge
Dai
Japanese
Kapla
Nair
Iyer
Iyangar
Coorghi
Bunt
Mbuti
1.00
0.75
0.50
0.25
0.00
from.y
0.00
0.03
0.06
0.09
drift parameter
migration weight
0.4
0.3
0.2
36.3 SE
-36.3 SE

H) m=7

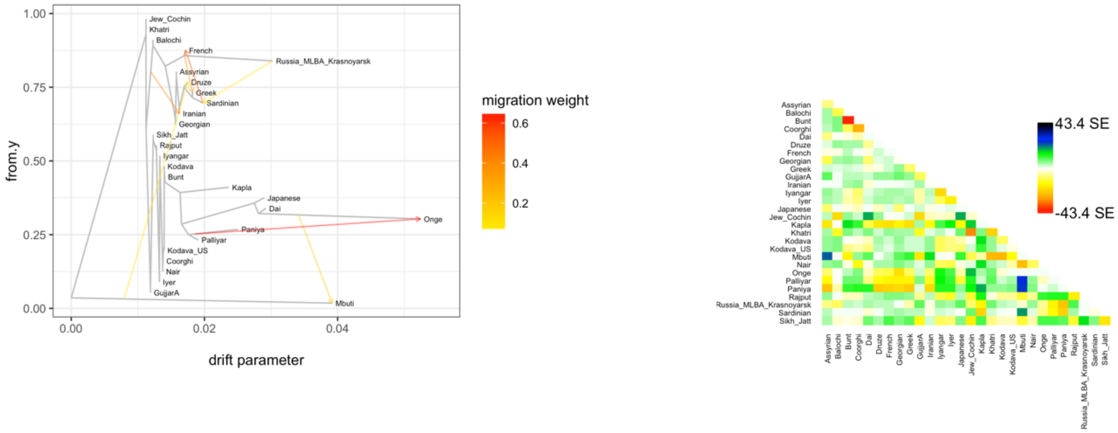
Jew_Cochin
Khatri
Balochi
French
Russia_MLBA_Krasnoyarsk
Assyrian
Druze
Greek
Sardinian
Iranian
Georgian
Sikh_Jatt
Rajput
Iyangar
Kodava
Bunt
Kapla
Japanese
Dai
Onge
Paniya
Palliyar
Kodava_US
Coorghi
Nair
Iyer
GujjarA
Mbuti
1.00
0.75
0.50
0.25
0.00
from.y
0.00
0.02
0.04
drift parameter
migration weight
0.6
0.4
0.2
43.4 SE
-43.4 SE

I) m=8

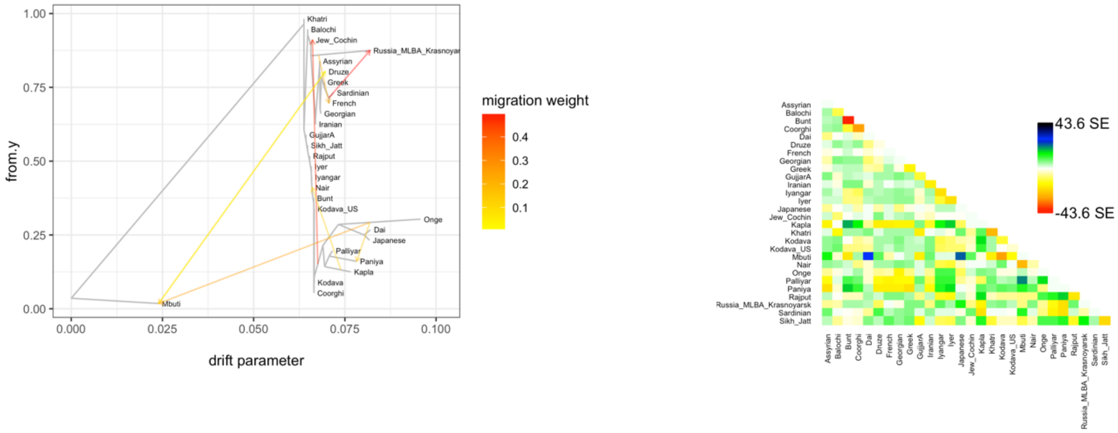
Khatri
Balochi
Jew_Cochin
Russia_MLBA_Krasnoyar
Assyrian
Druze
Greek
Sardinian
French
Georgian
Iranian
GujjarA
Sikh_Jatt
Rajput
Iyer
Iyangar
Nair
Bunt
Kodava_US
Onge
Dai
Japanese
Palliyar
Paniya
Kapla
Kodava
Coorghi
Mbuti
1.00
0.75
0.50
0.25
0.00
from.y
0.000
0.025
0.050
0.075
0.100
drift parameter
migration weight
0.4
0.3
0.2
0.1
43.6 SE
-43.6 SE

J) m=9

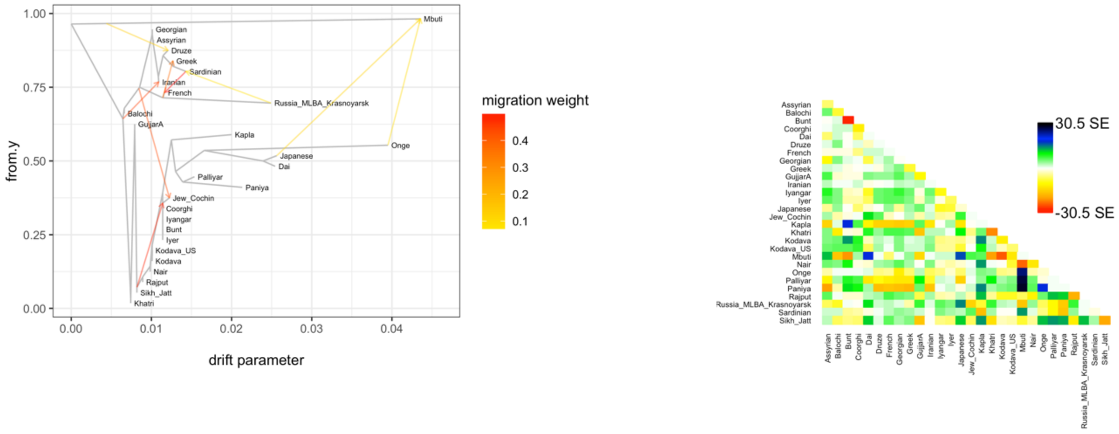

from.y
drift parameter
migration weight
0.4
0.3
0.2
0.1
Mbuti
Georgian
Assyrian
Druze
Greek
Sardinian
Iranian
French
Russia_MLBA_Krasnoyarsk
Balochi
GujjarA
Kapla
Onge
Japanese
Dai
Palliyar
Paniya
Jew_Cochin
Coorghi
Iyangar
Bunt
Iyer
Kodava_US
Kodava
Nair
Rajput
Sikh_Jatt
Khatri
30.5 SE
-30.5 SE


K) m=10

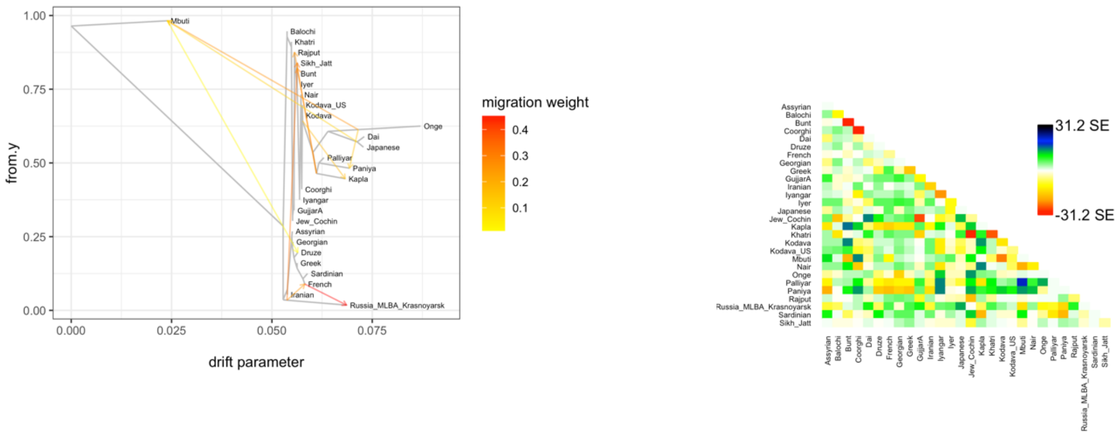

from.y
drift parameter
migration weight
0.4
0.3
0.2
0.1
Mbuti
Balochi
Khatri
Rajput
Sikh_Jatt
Bunt
Iyer
Nair
Kodava_US
Kodava
Onge
Dai
Japanese
Palliyar
Paniya
Kapla
Coorghi
Iyangar
GujjarA
Jew_Cochin
Assyrian
Georgian
Druze
Greek
Sardinian
French
Iranian
Russia_MLBA_Krasnoyarsk
31.2 SE
-31.2 SE

**Figure S10:** ***f*$_4$*-ratio* results.** Estimation of the proportion of ANI-related genetic ancestry in South Asians using Central Steppe MLBA (α).

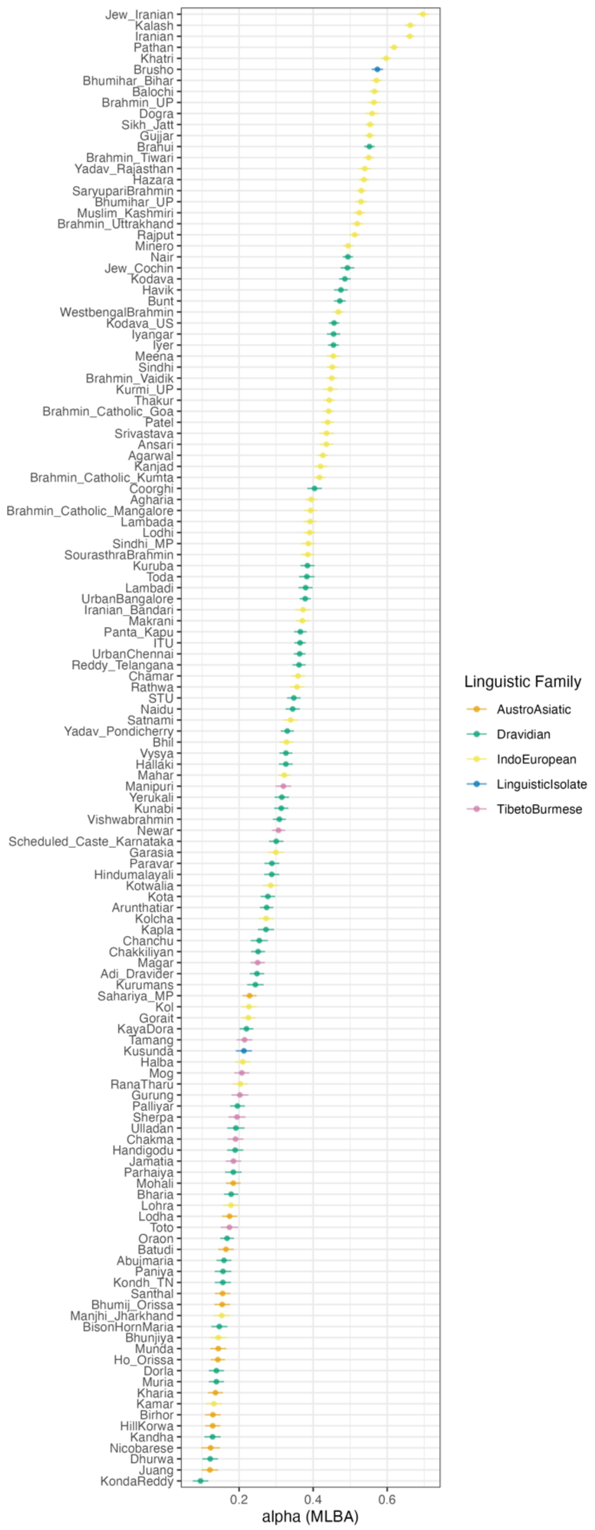

**Figure S11: D-statistic with source populations from qpAdm model.** D-statistic of the form *D*(South Asia, H2; Source; Mbuti), where H2 is one of the study populations and Source are either Central Steppe MLBA (A), Indus Periphery (B) or Onge (C).

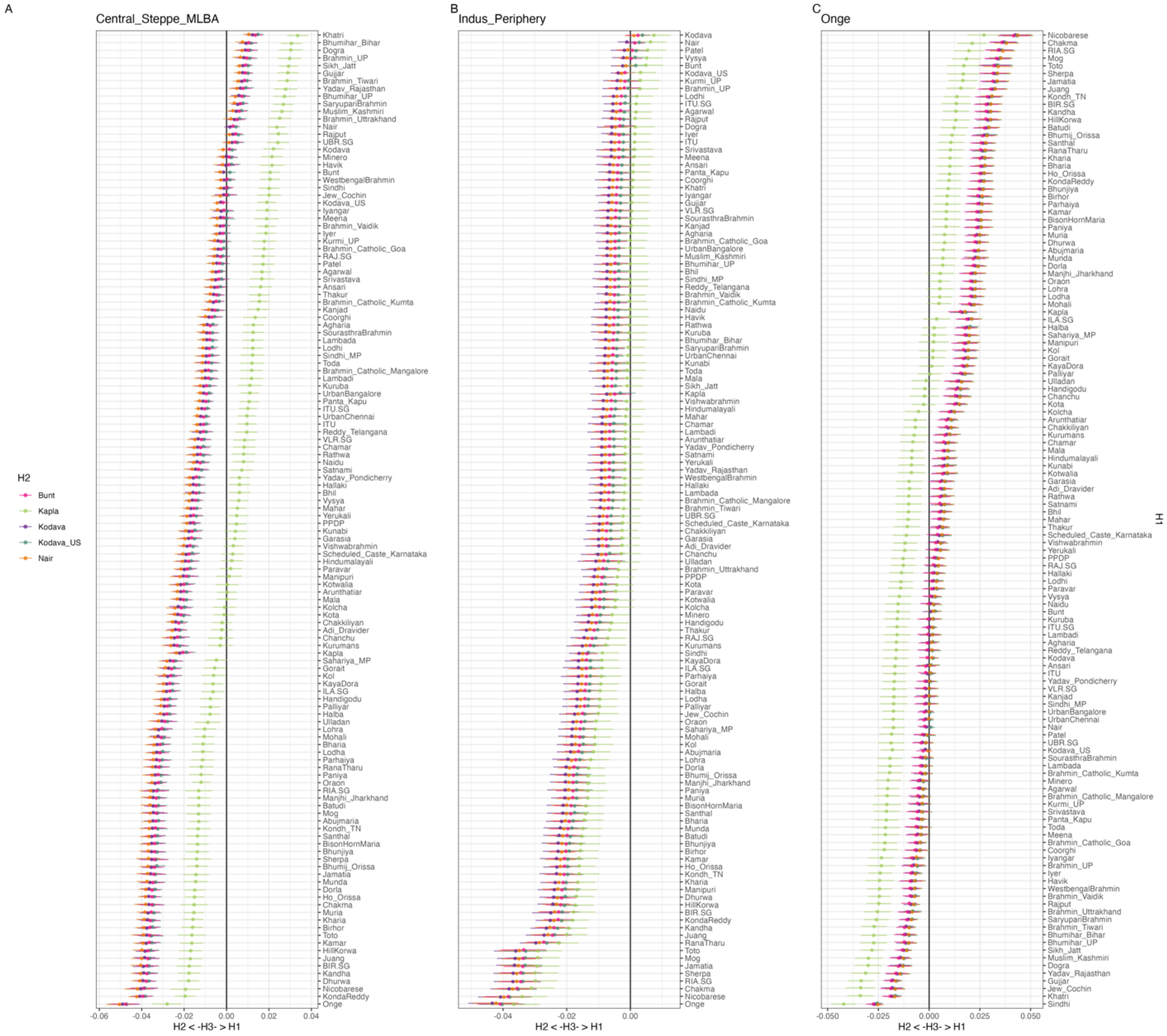

**Figure S12: D-statistic with source populations from qpAdm model using pseudo-haploid calls.** D-statistic of the form *D*(South Asia, H2; Source; Mbuti), where H2 is one of the study populations and Source are either Central Steppe MLBA (A), Indus Periphery (B) or Onge (C).

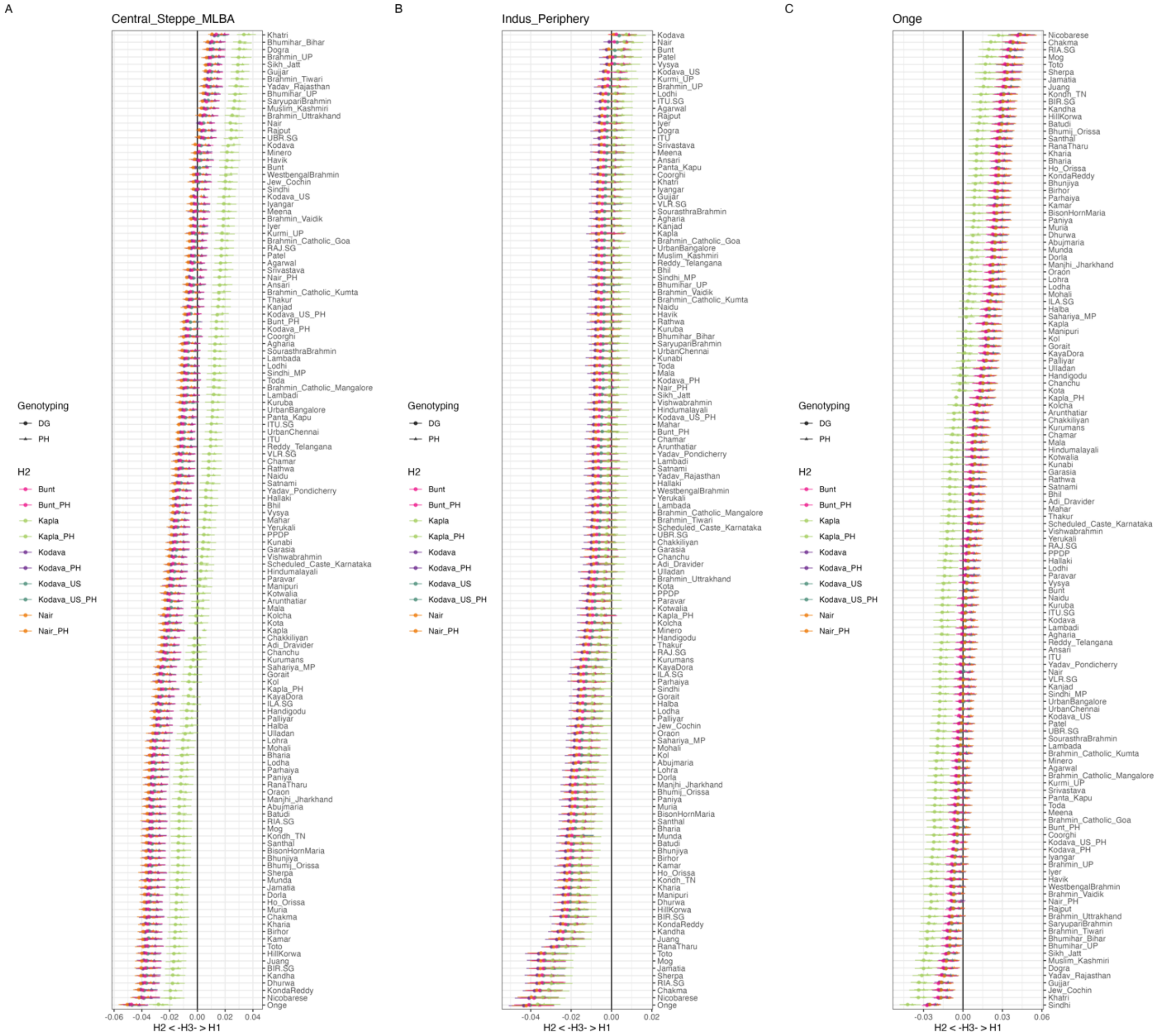

**Figure S13: Summary of co-ancestry matrix as average length count per population from ChromoPainter.**

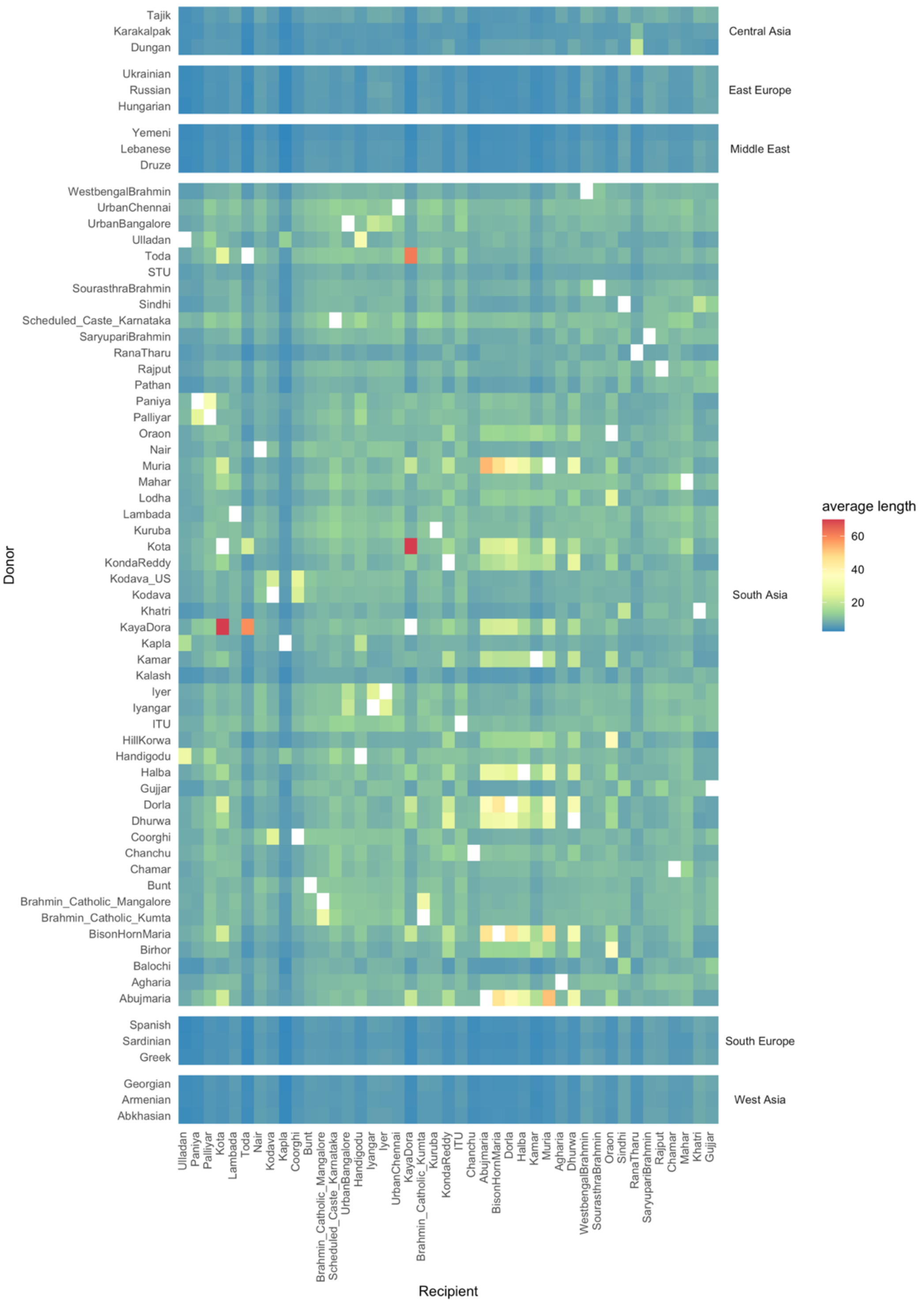

**Figure S14: fineSTRUCTURE clustering dendrogram.** This figure shows individual-level relatedness based on haplotypic similarity.

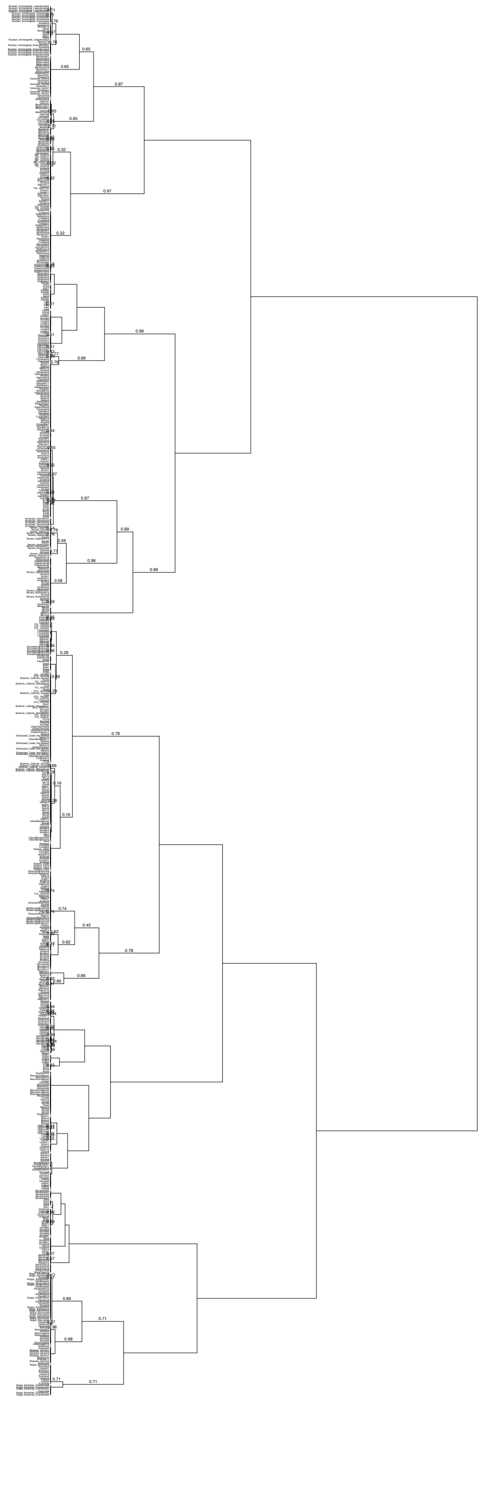

**Figure S15: Ancestry-specific haplotype copying across sampled populations from Southwest India.** The median per-locus copying probability from ChromoPainter represents the haplotype-level similarity between individuals from the focal populations from India – Rajput and Paniya added to reflect high ANI & high ASI populations respectively – and potential sources of Western Eurasian ancestry.

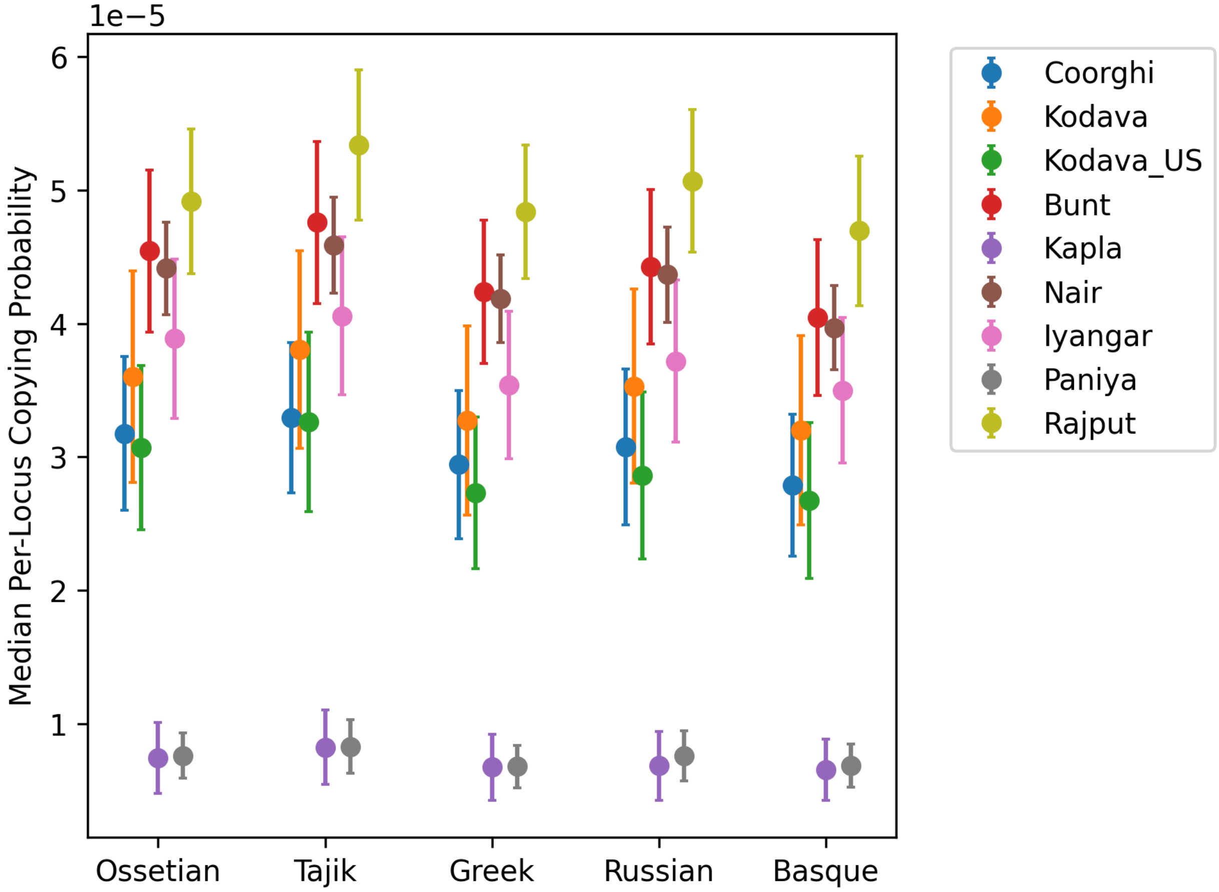

**Figure S16: Total length of ROH (SROH) versus the total number of ROH (NROH).** Averaged within populations.

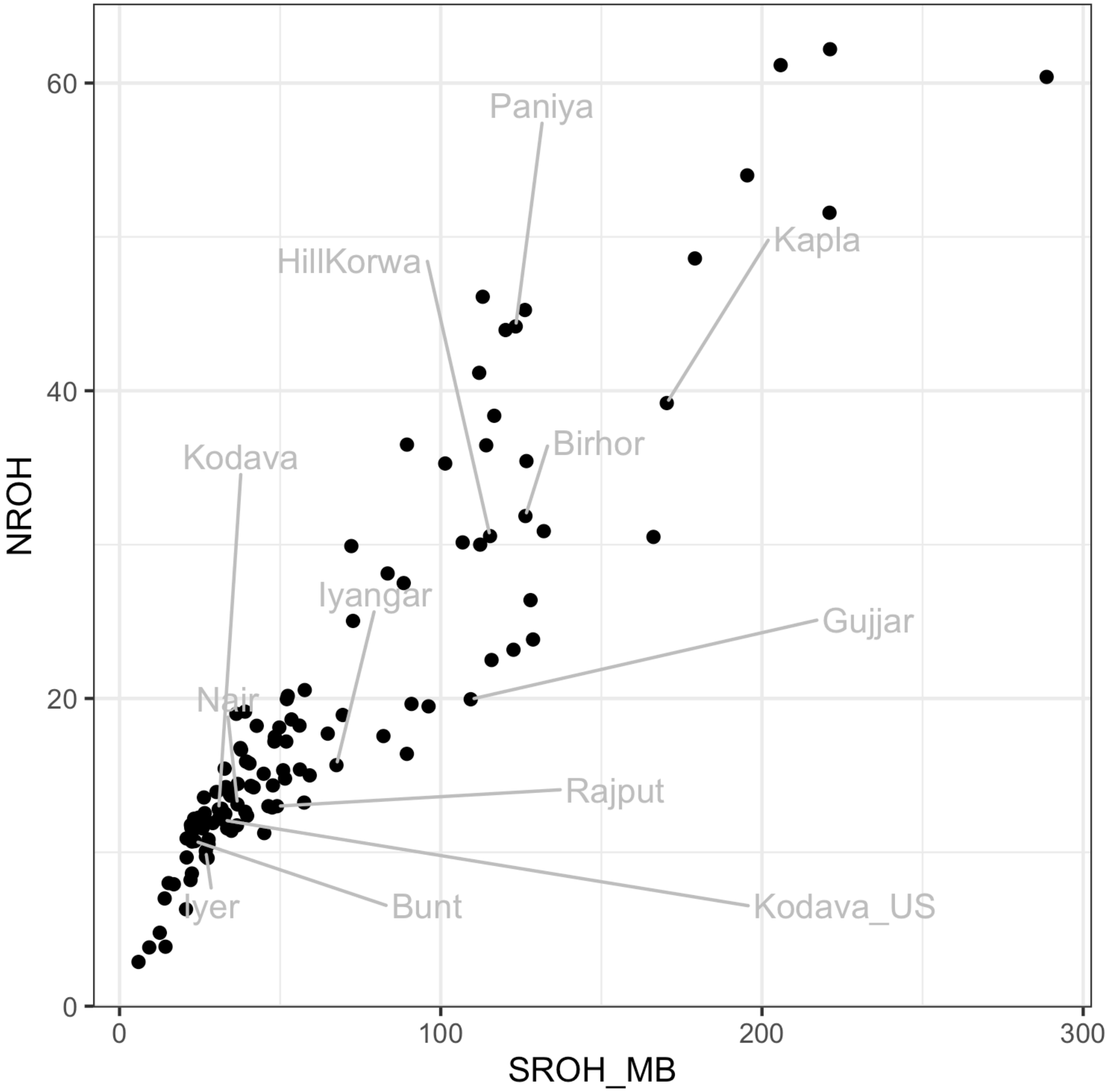

**Figure S17: Short versus long ROH for select populations.** A) Sum of total length of short (2-5Mb) ROH; B) Sum of total length of long (>10Mb) ROH.

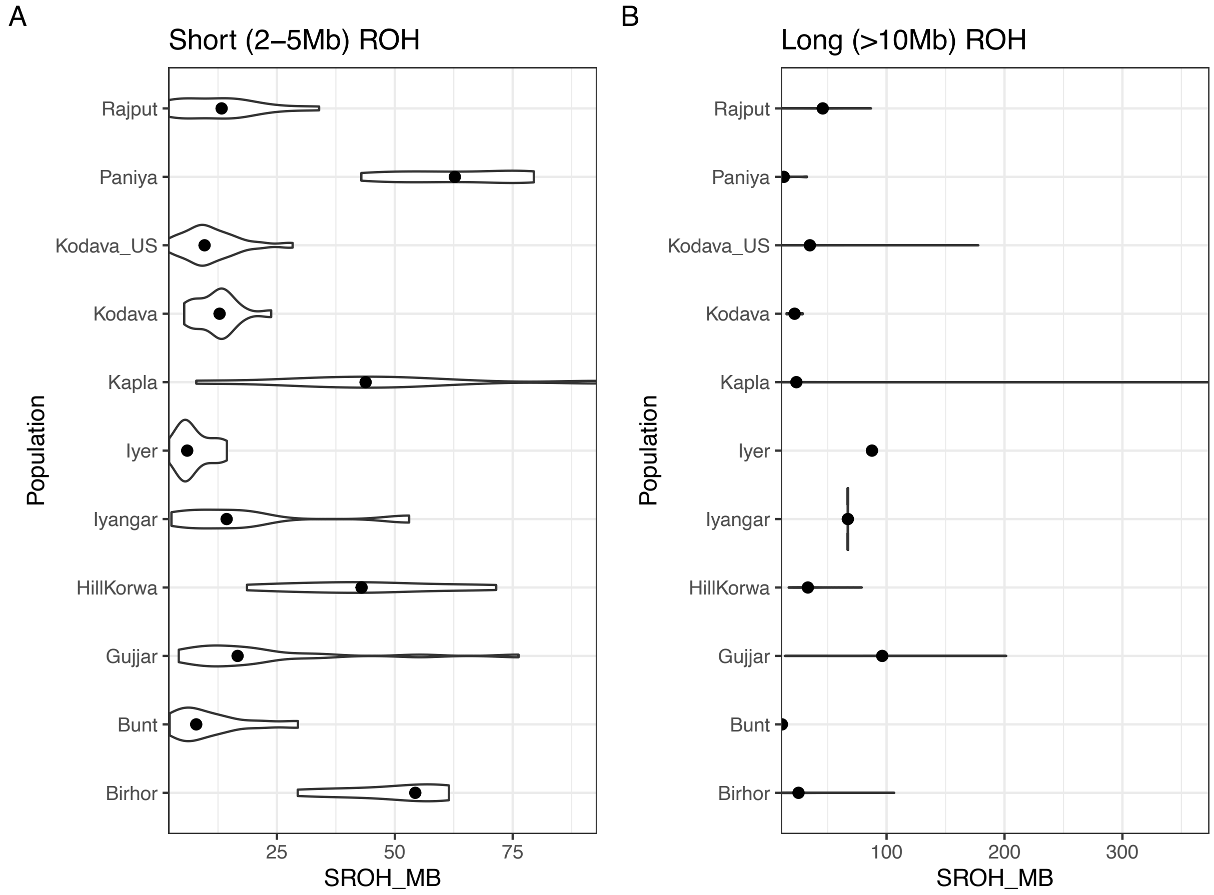

**Figure S18: Decay curves for the Southwest India populations (Kapla, Bunt, Kodava, Kodava US, and Nair) and other South Asian populations.** The X-axis represents the genetic distance in cM and the Y-axis the allele sharing correlation. The legend for each figure shows the mean and CI for the founder event (Tf) in generations before the present (gBP, 1 generation = 28 years), the intensity of the founder event (If), and the normalized root-mean-square deviation (NRMSD). The algorithm was run using Mbuti as the outgroup population since South Asian populations do not share a recent founder event with this African population. Populations names highlighted in yellow lacked evidence for a significant founder event (see Methods).

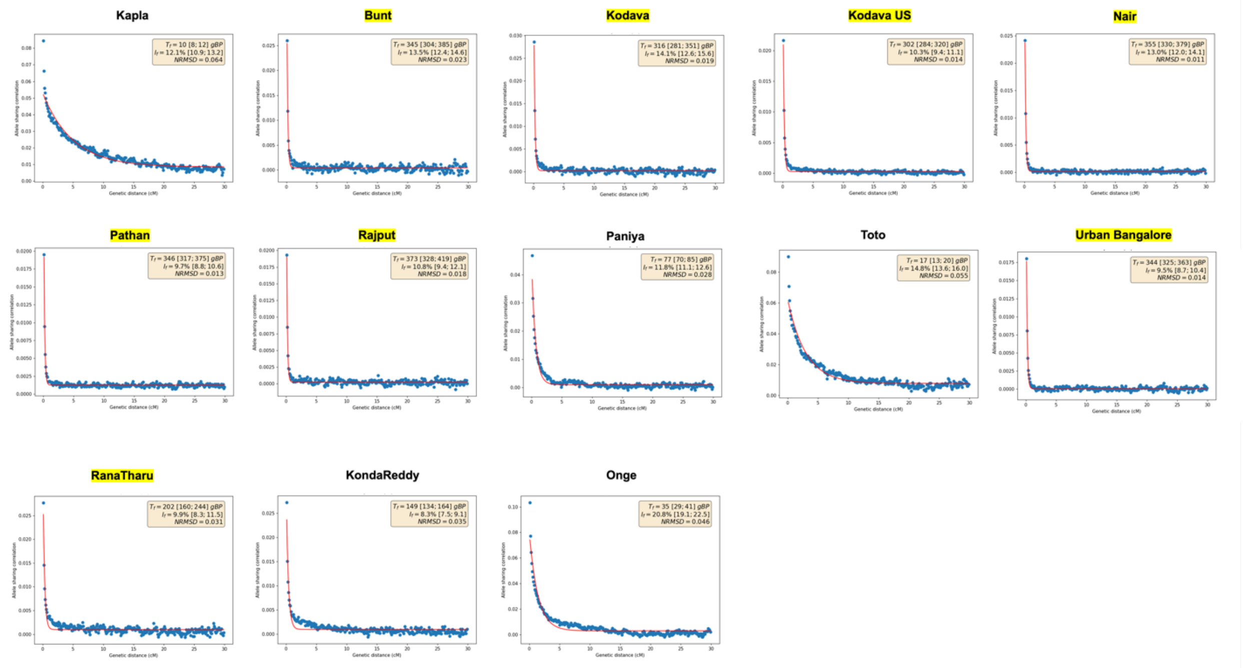

**Figure S19: Functional and novel variation in high-coverage WGS samples from Southwest India.** (A) Estimation of burden of functional mutations across all four study populations using 5000 binomial resampling iterations from carrier frequencies at functional alleles; (B) Estimating fraction of novel mutations per study population using binomial resampling of frequencies.

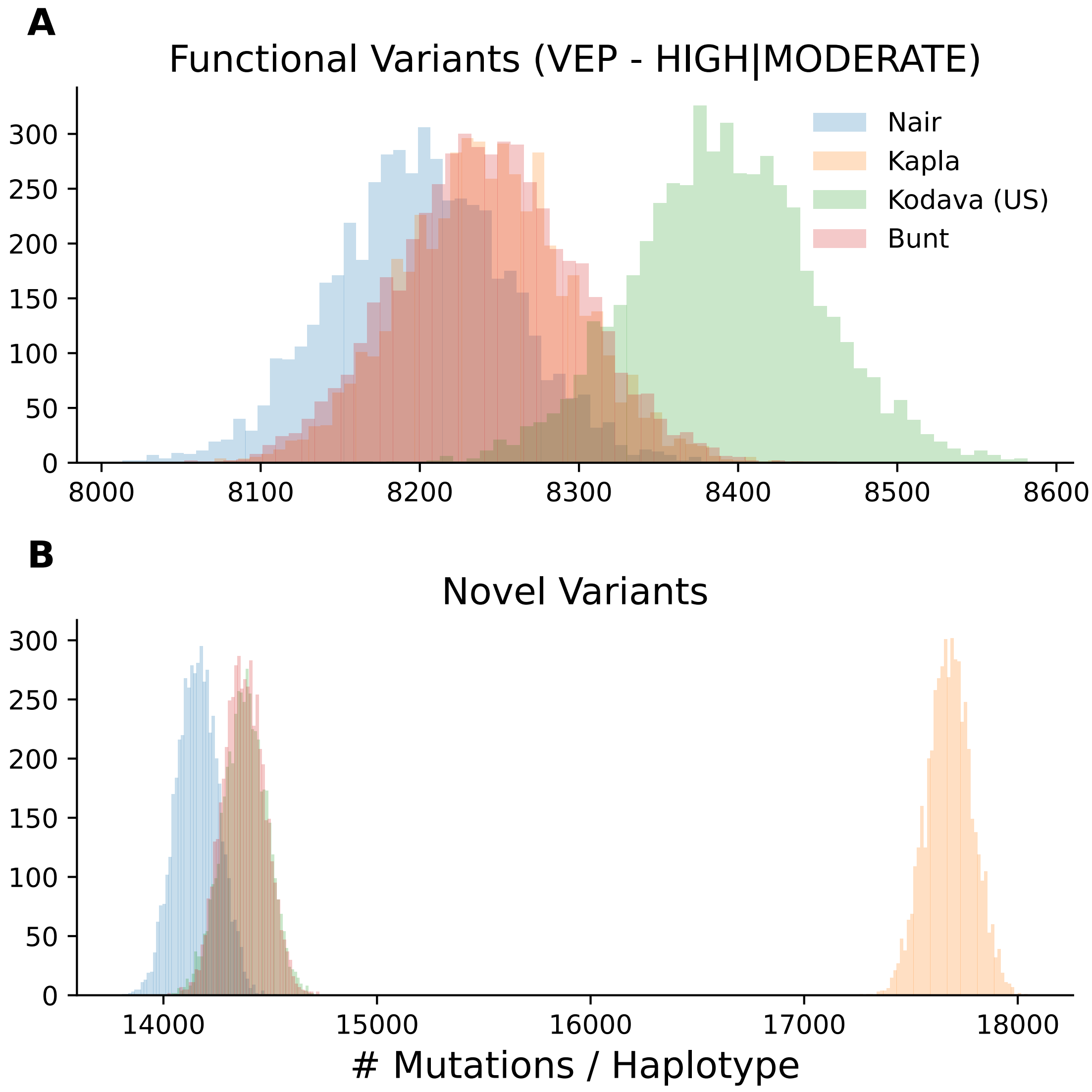

**Table S2: Populations included in Treemix analysis**

| Population Label | Sample size |
|---|---|
| Assyrian | 11 |
| Balochi | 21 |
| Bunt | 11 |
| Coorghi | 5 |
| Dai | 9 |
| Druze | 39 |
| French | 27 |
| Georgian | 13 |
| Greek | 20 |
| GujjarA | 19 |
| Iranian | 30 |
| Iyangar | 6 |
| Iyer | 13 |
| Japanese | 30 |
| Jew_Cochin | 5 |
| Kapla | 10 |
| Khatri | 11 |
| Kodava | 15 |
| Kodava_US | 78 |
| Mbuti | 10 |
| Nair | 44 |
| Onge | 6 |
| Palliyar | 11 |
| Paniya | 11 |
| Rajput | 14 |
| Russia_MLBA_Krasnoyarsk | 16 |
| Sardinian | 28 |
| Sikh_Jatt | 44 |

**Table S4: Populations used for calculating PCA-based SNP-loadings**

| Population Label | Sample size | Language Family |
|---|---|---|
| Handigodu | 6 | Dravidian |
| Toda | 12 | Dravidian |
| Paniya | 11 | Dravidian |
| Yadav_Pondicherry | 12 | Dravidian |
| Iyangar | 6 | Dravidian |
| Vysya | 39 | Dravidian |
| Iyer | 13 | Dravidian |
| Yerukali | 6 | Dravidian |
| Palliyar | 11 | Dravidian |
| UrbanChennai | 34 | Dravidian |
| STU | 8 | Dravidian |
| Arunthatiar | 18 | Dravidian |
| **Nair** | 44 | Dravidian |
| **Kodava_US** | 78 | Dravidian |
| Ulladan | 6 | Dravidian |
| UrbanBangalore | 34 | Dravidian |
| **Kodava** | 15 | Dravidian |
| ITU | 25 | Dravidian |
| **Bunt** | 11 | Dravidian |
| Chakkiliyan | 9 | Dravidian |
| Paravar | 7 | Dravidian |
| Panta_Kapu | 15 | Dravidian |
| **Kapla_A** | 6 | Dravidian |
| Adi_Dravider | 7 | Dravidian |
| SourasthraBrahmin | 9 | Indo-European |
| Agarwal | 36 | Indo-European |
| SaryupariBrahmin | 13 | Indo-European |
| Brahmin_Catholic_Goa | 14 | Indo-European |

| | | |
|---|---|---|
| Brahmin_Vaidik | 25 | Indo-European |
| Patel | 7 | Indo-European |
| Mahar | 19 | Indo-European |
| Brahmin_Catholic_Mangalore | 7 | Indo-European |
| Brahmin_Catholic_Kumta | 10 | Indo-European |
| Sindhi | 31 | Indo-European |
| Lambada | 11 | Indo-European |
| Brahmin_Tiwari | 15 | Indo-European |
| WestbengalBrahmin | 10 | Indo-European |
| Pathan | 37 | Indo-European |
| Khatri | 14 | Indo-European |
| Gujjar | 23 | Indo-European |
| Bhumihar_Bihar | 7 | Indo-European |
| Rajput | 17 | Indo-European |
| Sikh_Jatt | 44 | Indo-European |
| Bhil | 8 | Indo-European |
| Basque | 33 | Indo-European |

**Table S10: The ANI-ASI admixture timings in select populations from India inferred using ALDER (fit started at d > 0.30 cM). Study populations that were sequenced in this study are highlighted in bold. Admixture times inferred by ALDER are shown both in generations as well as in years (assuming a generation time of 28 years per generation (Narasimhan et.al.2019)). Language-family assignments are taken from (Nakatsuka et al. 2017; GenomeAsia100K Consortium 2019). Asterisks indicate admixture times whose decay rates have a Z-score >= 2 (corresponding to p-value < 0.05).**

| Population Label | Sample size | Language family | Admixture time (generations) | Admixture time SE (generations) | Admixture time (years) | Admixture time SE (years) |
|---|---|---|---|---|---|---|
| Handigodu | 6 | Dravidian | 94.69* | 16.76 | 2651.32 | 469.28 |
| Toda | 12 | Dravidian | 152.03* | 15.63 | 4256.84 | 437.64 |
| Paniya | 11 | Dravidian | 196.36* | 48.6 | 5498.08 | 1360.8 |
| Yadav_Pondicherry | 12 | Dravidian | 113.57* | 20.63 | 3179.96 | 577.64 |
| Iyangar | 6 | Dravidian | 130.8* | 22.13 | 3662.4 | 619.64 |
| Vysya | 39 | Dravidian | 150.91* | 11.95 | 4225.48 | 334.6 |
| Iyer | 13 | Dravidian | 110.54* | 8.94 | 3095.12 | 250.32 |
| Yerukali | 6 | Dravidian | 97.74* | 20.91 | 2736.72 | 585.48 |
| Palliyar | 11 | Dravidian | 131.9* | 28.8 | 3693.2 | 806.4 |
| UrbanChennai | 34 | Dravidian | 92.34* | 6.64 | 2585.52 | 185.92 |
| STU | 8 | Dravidian | 98.54* | 23.65 | 2759.12 | 662.2 |
| Arunthatiar | 18 | Dravidian | 138.24* | 15.31 | 3870.72 | 428.68 |
| **Nair** | 44 | Dravidian | 101.35* | 9.97 | 2837.8 | 279.16 |
| **Kodava_US** | 78 | Dravidian | 111.23* | 9.29 | 3114.44 | 260.12 |
| Ulladan | 6 | Dravidian | 153.22* | 37.22 | 4290.16 | 1042.16 |
| UrbanBangalore | 34 | Dravidian | 95.18* | 6.15 | 2665.04 | 172.2 |
| **Kodava** | 15 | Dravidian | 94.79* | 6.68 | 2654.12 | 187.04 |
| ITU | 25 | Dravidian | 109.87* | 9.92 | 3076.36 | 277.76 |
| **Bunt** | 11 | Dravidian | 107.83* | 9.05 | 3019.24 | 253.4 |
| Chakkiliyan | 9 | Dravidian | 118.73* | 26.87 | 3324.44 | 752.36 |
| Paravar | 7 | Dravidian | 104.17* | 14.98 | 2916.76 | 419.44 |
| Panta_Kapu | 15 | Dravidian | 109.2* | 19.69 | 3057.6 | 551.32 |
| **Kapla_A** | 6 | Dravidian | 112.65 | 173.92 | 3154.2 | 4869.76 |
| Adi_Dravider | 7 | Dravidian | 130.77* | 17.94 | 3661.56 | 502.32 |
| SourasthraBrahmin | 9 | Indo-European | 110.34* | 13.96 | 3089.52 | 390.88 |
| Agarwal | 36 | Indo-European | 110.58* | 6.69 | 3096.24 | 187.32 |
| SaryupariBrahmin | 13 | Indo-European | 111.85* | 9.87 | 3131.8 | 276.36 |
| Brahmin_Catholic_Goa | 14 | Indo-European | 94.11* | 9.58 | 2635.08 | 268.24 |
| Brahmin_Vaidik | 25 | Indo-European | 106.37* | 7.38 | 2978.36 | 206.64 |
| Patel | 7 | Indo-European | 95.14* | 18.16 | 2663.92 | 508.48 |
| Mahar | 19 | Indo-European | 116.8* | 13.37 | 3270.4 | 374.36 |
| Brahmin_Catholic_Mangalore | 7 | Indo-European | 100.66* | 13.77 | 2818.48 | 385.56 |
| Brahmin_Catholic_Kumta | 10 | Indo-European | 102.2* | 14.4 | 2861.6 | 403.2 |
| Sindhi | 31 | Indo-European | 94.68* | 7.01 | 2651.04 | 196.28 |
| Lambada | 11 | Indo-European | 87.3* | 17.35 | 2444.4 | 485.8 |
| Brahmin_Tiwari | 15 | Indo-European | 108.63* | 11.08 | 3041.64 | 310.24 |
| WestbengalBrahmin | 10 | Indo-European | 106.72* | 9.8 | 2988.16 | 274.4 |
| Pathan | 37 | Indo-European | 67.12* | 8.35 | 1879.36 | 233.8 |
| Khatri | 14 | Indo-European | 112.39* | 17.41 | 3146.92 | 487.48 |
| Gujjar | 23 | Indo-European | 93.09* | 7.3 | 2606.52 | 204.4 |
| Bhumihar_Bihar | 7 | Indo-European | 81.61* | 12.97 | 2285.08 | 363.16 |
| Rajput | 17 | Indo-European | 89.24* | 12.97 | 2498.72 | 363.16 |
| Sikh_Jatt | 44 | Indo-European | 60.4* | 5.82 | 1691.2 | 162.96 |
| Bhil | 8 | Indo-European | 65.68* | 10.01 | 1839.04 | 280.28 |

**Table S14: Haplotype and nucleotide diversity on mtDNA across matrilocal and patrilocal groups in Southwest India.** For each study population with available mitogenome data, we computed the number of unique haplotypes (h) and haplotype diversity and its standard error based on 100 bootstrap resampling iterations (HD, HD_SD) and the overall nucleotide diversity as a per-basepair rate (pi) using the software pixy (Korunes & Samuk 2021). See Methods for further details on calculations.

| Population | N | h | HD | HD_SD | pi |
|---|---|---|---|---|---|
| Bunt | 11 | 6 | 0.8363636364 | 0.0739667113 | 0.002278427863 |
| Kapla | 10 | 2 | 0.5333333333 | 0.2761616095 | 0.001332658228 |
| Kodava | 15 | 8 | 0.9238095238 | 0.04872801557 | 0.002026555868 |
| Kodava_US | 105 | 62 | 0.9873626374 | 0.002903768035 | 0.001853267642 |
| Nair | 44 | 34 | 0.9778012685 | 0.01163167342 | 0.002128760534 |
